# Multitrait engineering of Hassawi red rice for sustainable cultivation

**DOI:** 10.1101/2023.11.28.569140

**Authors:** Khalid Sedeek, Nahed Mohammed, Yong Zhou, Andrea Zuccolo, Krishnaveni Sanikommu, Sunitha Kantharajappa, Noor Al-Bader, Manal Tashkandi, Rod A. Wing, Magdy M. Mahfouz

**Affiliations:** Laboratory for Genome Engineering and Synthetic Biology, Division of Biological Sciences, King Abdullah University of Science and Technology (KAUST), Thuwal, Saudi Arabia; Center for Desert Agriculture, Biological and Environmental Sciences and Engineering Division (BESE), King Abdullah University of Science and Technology (KAUST), Thuwal, Saudi Arabia; Crop Science Research Center, Sant’Anna School of Advanced Studies, Piazza Martiri della Libertà 33, 56127 Pisa, Italy; Department of Biological Science, College of Science, University of Jeddah, Jeddah, Saudi Arabia; Arizona Genomics Institute, School of Plant Sciences, University of Arizona, Tucson, AZ, USA; International Rice Research Institute (IRRI), Strategic Innovation, Los Baños, 4031 Laguna, Philippines

**Keywords:** Hassawi rice, CRISPR, Sustainable agriculture, Trait engineering, Human nutrition, Genome sequencing, Metabolome screening

## Abstract

Sustainable agriculture requires locally adapted varieties that produce nutritious food with limited agricultural inputs. Genome engineering represents a viable approach to develop cultivars that fulfill these criteria. For example, the red Hassawi rice, a native landrace of Saudi Arabia, tolerates local drought and high-salinity conditions and produces grain with diverse health-promoting phytochemicals. However, Hassawi has a long growth cycle, high cultivation costs, low productivity, and susceptibility to lodging. Here, to improve these undesirable traits via genome editing, we established efficient regeneration and Agrobacterium-mediated transformation protocols for Hassawi. In addition, we generated the first high-quality reference genome and targeted the key flowering repressor gene, *Hd4*, thus shortening the plant’s lifecycle and height. Using CRISPR/Cas9 multiplexing, we simultaneously disrupted negative regulators of flowering time (*Hd2, Hd4*, and *Hd5*), grain size (*GS3*), grain number (*GN1a*), and plant height (*Sd1*). The resulting homozygous mutant lines flowered extremely early (∼56 days) and had shorter stems (approximately 107 cm), longer grains (by 5.1%), and more grains per plant (by 50.2%), thereby enhancing overall productivity. Furthermore, the awns of grains were 86.4% shorter compared to unedited plants. Moreover, the modified rice grain displayed improved nutritional attributes. As a result, the modified Hassawi rice combines several desirable traits that can incentivize large-scale cultivation and reduce malnutrition.

## 1. Introduction

Although rice (*Oryza sativa* L.) is a major staple food crop for caloric intake across the globe, the vast majority of health-promoting nutrients that are concentrated in the bran layer and embryo are removed via both the milling and polishing processes [1–4]. By contrast, “red rice” is generally consumed as a whole grain and contains higher concentrations of phenolic compounds, lipids, proteins, vitamins, fiber, amino acids, minerals, and other health-promoting compounds than white rice [4–8]. These compounds improve human health, especially phenolics and antioxidants, which possess anticancer, antidiabetic, and anti-inflammatory properties [5, 9, 10]. Consuming pigmented rice has been shown to enhance the immune system and reduce the risk of many chronic diseases and cancer [11].

The red pigmented rice – Hassawi - is a landrace indigenous to the Al-Ahsa region of the Eastern Province of Saudi Arabia, where it has been cultivated and consumed for hundreds of years as a whole grain, without milling or polishing [12, 13]. Our recent study of the metabolite and metal ion profiles of 63 Asian pigmented rice cultivars, including several Hassawi accessions, showed that Hassawi was the most nutritious red rice cultivar [4]. Hassawi rice is rich in antioxidants (proanthocyanidins), vitamins (B2, B6, C, and E), and essential minerals (zinc, iron, calcium, selenium, and strontium) [4]. Hassawi has high levels of ash and fat and a low glycemic load, which helps prevent type 2 diabetes [13, 14]. In addition, Hassawi rice exhibits remarkable adaptation to the local Saudi Arabian climate, showing resilience to high temperatures, drought, and soil salinity [15]. Despite these favorable attributes, farmers are reluctant to cultivate it due to its undesirable agronomic characteristics that include: 1) a lengthy growth cycle of ∼180 days (resulting in increased cultivation costs, water, and labor requirements); 2) a plant height of approximately 185 cm (i.e. ∼6 ft, making them prone to lodging which can cause significant yield losses) [16]; and, 3) is low yielding (i.e. ∼ 2 tons/ha) as compared to other high yielding varieties (i.e. 5-10 tons/ha). Improvement of these three key traits *via* genome engineering has the potential to produce a 21^st^ century rice for the KSA and the Middle East that is not only highly nutritious but would also have a shorter life cycle, be lodging resistant and high-yielding.

Rice is a facultative short day (SD) plant whose flowering is promoted under short day (SD) conditions (<10-h light/day) and inhibited under long day (LD) conditions (>14-h light/day) [17]. Heading date in rice is a quantitative trait controlled by several genetic and environmental factors. Extensive research has identified several photoperiod flowering time suppressors in the Early heading date 1 (Ehd1) pathway of rice, including *Hd2*, *Hd4,* and *Hd5* [18–22]. Disrupting these genes can promote early heading date and reduce the rice plant height [4]. Plant yield is a key agronomic trait that is determined by many components, including the number of panicles per plant, number of grains per panicle, and grain size and weight [23, 24]. Several studies have identified a number of yield-related genes controlling these traits in rice and other crops. For example, *GRAIN SIZE 3* (*GS3*), and *GRAIN NUMBER 1a* (*Gn1a*) regulate grain size and grain number, respectively, in rice [25–27] and are excellent targets for genome editing. Finally, shorter cultivars are usually preferred by farmers and breeders, as they are resistant to lodging, consume fewer soil nutrients, and require less fertilizer to grow and flourish. Dwarf IR8 rice contains a natural mutation of *Semi-dwarf 1* (*Sd1*), encoding a GA20 oxidase 2 (GA20ox-1) that functions in gibberellin biosynthesis [28]. Therefore, *Sd1* is considered to be a good target for CRISPR/Cas9 mutagenesis to enhance the lodging resistance of plants [29].

In this study, we used the CRISPR/Cas9 genome editing technology to improve Hassawi rice by shortening the growth cycle, reducing its susceptibility to lodging, and improving its yield by first establishing an efficient regeneration/Agrobacterium-mediated transformation system for the delivery of the CRISPR materials into rice cells. Next, we generated a high-quality reference genome of Hassawi rice to accurately identify target loci and design the necessary guide RNAs for genome editing. Finally, we employed the CRISPR/Cas9 system to knock out the *Hd4* gene, a negative regulator of flowering time, and to simultaneously knock out six genes that negatively influence flowering time (*Hd2, Hd4*, and *Hd5*), grain size and number (*GS3 and Gn1a*), and plant height (*Sd1*), which resulted in the creation of several Hassawi rice lines that exhibited remarkably fast growth rates, increased yield, and improved lodging resistance. Finally, we assessed the notational quality of the modified rice grain by screening its metabolome using the Ultrahigh Performance Liquid Chromatography-Tandem Mass Spectroscopy (UPLC-MS/MS).

## 2. Materials and Methods

### 2.1. Plant material and explant preparation

Hassawi rice (accession 190) was obtained from the herbarium of the King Abdullah University of Science and Technology and was originally collected from Habib Al-Attar farm in the Al Ahsa region, Saudi Arabia. Mature dry seeds were used as explants for callus induction. The seeds were dehusked and sterilized with 70% (v/v) ethanol for 1 min and 30% (v/v) commercial bleach (5.25% sodium hypochlorite) solution containing one drop of Tween 20 for 45 min with continuous shaking. The seeds were rinsed five times with sterile distilled water and dried on Whatman paper for 5 min before being used for callus induction.

### 2.2. Embryogenic callus induction and regeneration

For callus induction, we tested the modified 2NBK medium, which we previously used to induce calli from black rice [4]. This medium contains 2 mg/l of the synthetic auxin 2,4-D; the detailed compositions of all media are described in Supplementary Table S1. Thirty-six seeds were placed on callus-induction medium with the scutella facing up and incubated at 32°C under continuous light for 7 days. The scutella were separated from the seeds and subcultured in fresh 2NBK medium for another 7 days under the same conditions. The embryogenic calli were induced by subculturing the calli on nNBKC for seven days at 32°C under continuous light. The calli were subcultured on R8 regeneration medium, which was developed for shoot induction in black rice [4]. The regeneration frequency was calculated based on the number of regenerated shoots compared to the total number of calli. To induce root development, the regenerated shoots were transferred to Magenta boxes containing MSRO rooting medium under the same temperature and light conditions.

### 2.3. Agrobacterium-mediated transformation of Hassawi rice

To establish a transformation protocol for Hassawi rice, we used the binary vector pRGEB32. This vector contains the *Hygromycin phosphotransferase* (*Hpt*) gene in the T-DNA region to enable the selection of transformed cells. The vector was electroporated into *Agrobacterium tumefaciens* strain EHA105 using a Bio-Rad Laboratories pulser. The bacterial cells were cultured on an AB plate containing 34 mg/L chloramphenicol and 50 mg/L kanamycin in the dark at 28°C for 3 days according to Hiei and Komari (2008). The growing bacteria were collected with a loop and suspended in 1 mL of AAM medium at a final density OD_600_ = 0.2.

Calli produced from mature seeds were cut into small pieces (0.5–1 mm in diameter) using a sharp scalpel and forceps. The calli were cultivated on fresh 2NBK medium at 32°C under continuous light for 3 days. New, actively proliferating calli were collected and mixed with the agrobacterium suspension for 2 min at room temperature. The calli were dried well on sterilized Whatman paper before being placed on co-cultivation medium and incubated at 25°C in the dark for 3 days. After the co-cultivation step, the calli were placed on the first selection medium (NBKCH50) containing a high concentration of hygromycin (50 mg/L) and incubated for 14 days to select transformed cells. The surviving calli were transferred to the second selection medium (nNBKCH65) containing a higher concentration of hygromycin (65 mg/L) and incubated for seven days. The embryogenic calli were transferred to R8h65 medium containing 65 mg/L hygromycin and incubated for 14 days. The regenerated plantlets were transferred to Magenta boxes containing a suitable free-hormonal-rooting medium (MSROH65) supplemented with hygromycin (65 mg/L). All culture steps, from callus induction to rooting, were conducted at 32°C under continuous light, except for the co-cultivation step. When the seedlings were mature, with a well-developed root system and leaves touching the cap of the Magenta box, they were transferred to soil and grown under greenhouse conditions.

### 2.4. Validation of transformed plants

Approximately 0.5 g of fresh leaf tissue from the transformed plants was subjected to total DNA extraction using the DNAquick Plant System (Tiangen Biotech), according to the manufacturer’s protocol. DNA quality and concentration were verified by analysis in a NanoDrop spectrophotometer and 1% agarose gel electrophoresis. PCR was performed to amplify 240 bp of the T-DNA transgene using Phusion High-Fidelity DNA Polymerase (Thermo Fisher Scientific) and a pair of primers (Cas9_F7, and Nos_R7) (Supplementary Table S2). The PCR products were visualized on 1.2% agarose gels.

### 2.5. Plant material, library construction and genome sequencing

*O. sativa* cv. Hassawi (IRGC No.121913) seeds were obtained from IRRI and planted in a greenhouse at the University of Arizona, USA. Young, healthy and dark treated (>24hrs) leaf tissue was collected from a single seedling, flash-frozen in liquid nitrogen, and ground to a fine powder using a frozen mortar and pestle. The CTAB method [30], with slight modifications, was used to extract high molecular weight genomic DNA from the tissue. The quantity and quality of the genomic DNA were validated using a Qubit HS Fluorometer (Invitrogen, USA) and a NanoDrop spectrophotometer, respectively. The DNA was digested with the restriction enzymes *Eco*RI and *Hind*III to confirm accessibility to molecular reagents, and the size was validated by pulsed-field electrophoresis. The size of the genomic DNA was checked using a Femto Pulse System (Agilent). A 15 µg sample of DNA was sheared to an appropriate size range (10-30 kb) using a Covaris g-TUBE. Bead purification was performed using PB Beads (PacBio, Menlo Park, CA). A PacBio sequencing library was constructed following the manufacturer’s protocol using a SMRTbell Express Template Prep kit 2.0 and size selected on a BluePippin instrument (Sage Science) using the S1 marker with 10-25 kb size selection. The library was quantified using a Qubit HS kit (Invitrogen), and its size was checked on a Femto Pulse System (Agilent). The sequencing library was prepared for sequencing with a PacBio Sequel II Sequencing kit 2.0 for HiFi libraries, loaded onto a single 8M SMRT cell, and sequenced in CCS mode in a Sequel II instrument for 30 hours.

### 2.6. Hassawi de novo genome assembly

The Hassawi (IRGC No.121913) genome was assembled to a PSRefSeq quality level following the pipeline used for the 12 *O. sativa* genome data set described by [31]. A *de novo* genome assembly was performed using Hifiasm v 0.15.5. Genome Puzzle Master (GPM) [32] was used for chromosome scaffolding and gap fixing, and the genome sequence of rice accession MH63RS2 was used as a reference guide. Bionano optical maps were generated using high-molecular weight DNA at Corteva, Bionano Solve (v.3.4) was used for genome mapping, and Bionano Access (v.1.4) (Bionano Genomics; https://bionanogenomics.com/) was used for optical map analysis. The PacBio raw data and genome sequences were deposited at NCBI under BioProject ID: PRJNA657951.

### 2.7. Short-read sequencing of Hassawi (accession 190)

To validate the accession of Hassawi that was used for transformation, seeds from another Hassawi rice accession (accession 190) were germinated and leaf samples were collected to extract DNA and to perform short read sequencing (PE 2×150 bp) using the Illumina NovaSeq6000 platform. Short reads were checked for quality and trimmed for adaptors using Fastqc v 0.11.9 [33] and Trimmomatic v 0.38 [34], respectively. The short reads from Hassawi accession 190 were mapped onto our newly generated reference Hassawi genome. In addition, 10 more accessions (>14X) of each of 16 subpopulations from 3K-RGP plus Hassawi (a total of 161 accessions) were subjected to principal component analysis (PCA) based on genome-wide SNPs (missing rate 0.2, maf 0.01, and prune in --indep-pairwise 50 1 0.8) using Plink [35]. The SNPs were called using IRGSP as a reference following the high-performance HPC-GVCW workflow [36] based on GATK4 best practice pipeline (https://github.com/IBEXCluster/Rice-Variant-Calling).

### 2.8. Structural variation, SNP analysis, and Transposable element (TE) quantification and masking

Structural variation analysis was performed following standard workflows (RPFP, https://yongzhou2019.github.io/Rice-Population-Reference-Panel/software/) for a rice population reference panel [37]. Briefly, using the Hassawi genome as a reference, the presence and absence of SVs was analyzed using NGMLR [38] as an aligner and SVIM [39] as a caller. Inversions were called using nucmer [40] as an aligner and Synteny and Rearrangement Identifier (SyRI) [41] as the SV caller. SNP calling for the 3K Rice Genome Project (3K-RGP) [42] using Hassawi as a reference was performed following the GATK4 best practice pipeline (https://gatk.broadinstitute.org/hc/en-us/sections/360007226651-Best-Practices-Workflows). TE content in Hassawi was assessed using RepeatMasker [43] run with the TE library rice7.0.0.liban.txt [31]. The soft-masked genome assembly was then used for gene prediction.

### 2.9. RNA-seq and Iso-seq library preparation and sequencing for baseline genome annotation

RNA was extracted from Hassawi (IRGC No.121913) leaf and root tissue using a Maxwell RSC Plant RNA Kit and a Maxwell RSC48 instrument following Promega’s protocol. RNA samples were multiplexed in a NovaSeq 6000 S1 flow cell, and short read sequencing of total RNA was performed (PE 2×150 bp). The PacBio Iso-seq sequencing library was prepared following the standard protocol for isoform sequencing using a SMRTbell Express Template Prep Kit 2.0, after the cDNA synthesis step. cDNA samples from the two tissues were then barcoded and amplified. The multiplexed library was prepared for sequencing using a PacBio Sequel II Sequencing kit 2.0, loaded onto a single 8M SMRT cell, and sequenced in CCS mode in a PacBio Sequel II instrument for 30 hours.

### 2.10. De novo gene-coding annotation with long and short RNA reads

Raw paired-end (PE) short RNA-seq reads from leaf and root tissue were trimmed and filtered using sickle v 1.33 [44], retaining 596,399,126 (99%) and 466,217,065 (98%) of the raw reads, respectively. RNA-seq reads were then assembled using Trinity v. 2.8.4 [45] with default parameters, plus the read-normalization parameter. Transcripts were clustered using cd-hit v. 4.6.8 [46] with a minimum of 98% identity, and yielded 502,591 assembled transcripts reduced from 714,289. For the raw iso-seq reads, the isoseq3 v.3 pipeline [47] was utilized to filter and cluster raw reads into high-quality (HQ) non-chimeric full-length transcripts, yielding 182,759 HQ transcripts, with an average length of 2,304.58 nt.

For *de novo* gene prediction, the Gene Finding module was run using the genome assembly soft-masked for TEs and both transcriptomes as direct evidence with default settings in OmicsBox 2.2.4. [48]. The translated annotation was accessed using BUSCOv.3.0 [49] against the Viridiplantae database v.10. Lastly, the *de novo* annotated CDSs were aligned against the *O. sativa* IRSPG-1.0 annotated CDS using BLAST+ v 2.7.1 to assess completeness. GENESPACE was used to call putative orthologs and for synteny analysis of the Hassawi and Nipponbare genomes [50].

### 2.11. Design of sgRNAs, vector construction, and Agrobacterium-mediated transformation

To identify the sequences of the target genes in Hassawi rice, BLAST analysis against the sequenced Hassawi genome was performed using the Nipponbare *Hd2/DTH7* (Os07t0695100-01), *Hd4/Ghd7* (Os07t0261200-01), *Hd5/DTH8* (Os08t0174500-02), *GS3* (Os03g0407400), *Gn1a* (Os01g0197700), and *Sd1* (Os01g0883800) sequences as queries. One sgRNA was designed to target the second exon of *Hd4*, and the other sgRNAs were designed to target the first exons of the remaining target genes. Twenty nucleotides in front of the NGG protospacer-adjacent motif (PAM) sequence were selected using the CHOPCHOP design tool [51]. The sgRNAs oligos were synthesized by GenScript (www.genscript.com) as complementary fragments containing a four-nucleotide overhang for the BsaI site (oligos are presented in Supplementary Table. 2). The forward and reverse fragments were annealed in a PCR machine. The sgRNAs were cloned into the BsaI-digested pRGEB32 vector under the control of the *OsU3* promoter. The pRGEB32 vector contains *Cas9* driven by the *OsUbiquitin* promoter and a hygromycin resistance gene driven by the *35S* promoter.

For multiplex gene targeting, the endogenous tRNA-processing system that was previously shown to work in rice was utilized [52]. The tRNA–gRNA construct flanked by the BsaI restriction sites was synthesized by GenScript and ligated into the pRGEB32 vector. All seven binary vectors were transformed into TOP10 *E. coli* competent cells, and a few colonies were selected for plasmid extraction. The clones were digested by BsaI and sequenced by Sanger sequencing to confirm the correct insertion of the sgRNA sequences and the tRNA–gRNA construct before the RNA scaffold. The validated construct was introduced into *Agrobacterium tumefaciens* strain EHA105 by electroporation, and positive colonies were selected on LB agar plates containing 25 mg/L chloramphenicol and 50 mg/L kanamycin.

### 2.12. Rice transformation and detection of on-target mutations

Agrobacterium cells harboring pRGEB32 containing the sgRNA sequences were used to transform rice calli using our optimized protocol (described above). Genomic DNA was extracted from fresh leaves of the transformed plants using our laboratory-prepared extraction buffer following the protocol of Iowa State University. PCR amplification was carried out using T-DNA sequence-specific primers (Cas9_F7 and Nos_R7) to verify the integration of the CRISPR materials in the rice genome. PCR products were visualized by 1.2% agarose gel electrophoresis. Plants containing T-DNA were used to amplify the potential target site using a primer pair flanking the sgRNA binding site. The PCR amplicons were visualized and purified from the gel using a QIAquick Gel Extraction Kit (Qiagen) and cloned into a pJET vector using a CloneJET PCR Cloning Kit (Thermo Fisher Scientific). The pJET constructs were transformed into TOP10 *E. coli* competent cells, and the colonies were selected on kanamycin plates for plasmid extraction and Sanger sequencing. The presence of on-target mutations was determined by alignment against the wild-type sequence using SnapGene software (GSL Biotech; https://www.snapgene.com/).

### 2.13. Trait phenotyping

The edited plants and their wild-type controls were grown in the KAUST greenhouse at 30°C under natural sunlight. Heading date was calculated based on the date of seed germination to the emergence of the first panicle. Plant height was measured when the first panicle emerged using a ruler. Other phenotypic traits were measured when the plants reached full maturity. Tiller number per plant was counted, panicles were harvested, and the number of filled and unfilled spikelets per panicle was counted and used to calculate the seed setting rate (ratio of the number of filled grains to the total number of spikelets). The rough paddy rice grain length and width were measured with ImageJ software, and 1000 dehusked grains were weighed using a sensitive electronic balance (0.001 g sensitivity). The total yield per plant was measured as the total weight (grams) of grain collected per plant. All data were statistically analyzed by unpaired *t*-test using GraphPad Prism 9.3.1. Significant differences are indicated by asterisks: *, *p*<0.05; **, *p*<0.01; ***, *p*<0.001; ****, *p*<0.0001.

### 2.14. Untargeted metabolic profiling of Hassawi grain

Rice samples were collected from a pool of rice individuals, and three biological replicates were analyzed for each line (*hd4, Mx,* and WT). The dehusked grains were ground in liquid nitrogen and lyophilized for 30 h, after which the samples were prepared by Metabolon Inc. (Durham, NC, USA). Several recovery standards were added prior to the extraction process. The samples underwent extraction using 80% methanol with vigorous shaking for 2 min (Glen Mills GenoGrinder 2000). The extract was split into four fractions: two for positive ion mode electrospray ionization (ESI) RP/UPLC–MS/MS analysis, one for negative ion mode ESI RP/UPLC–MS/MS analysis, and one for negative ion mode ESI HILIC/UPLC–MS/MS analysis. All methods utilized a Waters ACQUITY UPLC and a Thermo Fisher Scientific Q-Exactive high-resolution/accurate mass spectrometer connected to a heated ESI (HESI-II) source and Orbitrap mass analyzer operated at 35,000 mass resolution. The extract was then dried and subsequently reconstituted in solvents compatible with each of the four methods, containing standardized concentrations of reference compounds to ensure consistent injection and chromatographic results. Raw UPLC–MS/MS data were filtered to eliminate system artifacts, misassignments, redundancy, and background noise. Compounds and peaks were identified by comparing them to a library of purified standards, and quantification was performed based on area-under-the-curve detector ion counts, with Welch’s two-sample t-tests applied for data analysis.

## 3. Results

### 3.1. Establishment of efficient regeneration protocol for Hassawi rice

The ability to introduce transgenes into plant cells and to regenerate these cells into healthy plants is a major bottleneck for the widespread use of genome editing for crop improvement [53]. As rice is a model monocot crop, much effort in the last decade has focused on optimizing its regeneration and transformation methods. However, this analysis has largely focused on the *japonica* variety Nipponbare, which is relatively easy to transform and regenerate. Other rice germplasm can vary significantly in their response to callus induction and their regeneration and transformation efficiencies [54]. To our knowledge, no efficient regeneration protocol is currently available for Hassawi rice. Therefore, we first aimed to develop an efficient regeneration protocol for Hassawi rice via somatic embryogenesis. We tested callus inducibility of Hassawi using our recently developed protocol for the Indonesian black rice cv. Cempo Ireng [4]. We initiated *in vitro* callus formation from mature grains using 2NBK medium (Supplementary Fig. S1A, media components are listed in Supplementary Table S1). A typical yellowish, friable callus mass emerged after 7 days of culture, with a callus induction frequency (CIF) of approximately 93.9% (Supplementary Fig. S1B). We separated the scutella from the seeds, transferred them to fresh medium, and incubated them under the same conditions for another 7 days to increase the tissue mass (Supplementary Fig. S1C). We then induced somatic embryogenesis by applying osmotic pressure for 7 days using medium containing a high concentration of sugars (nNBKC), including sorbitol (55 g/L) and sucrose (20 g/L). We also tested the effects of culture on the best regeneration medium developed for black rice: this medium contains Murashige and Skoog (MS) basal salts supplemented with 1 mg/l 1-naphthaleneacetic acid (NAA) + and 2 mg/l 6-benzylaminopurine (BAP). We did not observe any shoot induction during the first week of cultivation. However, when we transferred the calli to fresh regeneration medium, green shoots emerged after 3–7 days of culture, with a total frequency of approximately 21.8% (Supplementary Fig. S1D). The shoots developed good root systems on hormone-free medium (MSRO) and matured into healthy seedling in 7-10 days (Supplementary Fig. 1E). The seedlings were transferred to soil for acclimatization in the greenhouse (30°C, natural sunlight) (Supplementary Fig. S1F).

### 3.2. Establishment of agrobacterium transformation system for Hassawi rice

In an attempt to develop a highly efficient transformation system, we used *Agrobacterium tumefaciens* strain EHA105 harboring the binary vector pRGEB32, which contains the hygromycin resistance gene as a selectable marker. To test our optimized regeneration protocol for recovering transformed plants, we transfected the induced calli with the agrobacterium suspension and selected transformants by culturing on NBKCH50 medium containing 50 mg/L hygromycin for 14 days (Fig. 1B). We then induced somatic embryogenesis and cultured the surviving calli on nNBKCH65 medium containing 65 mg/L hygromycin for seven days (Fig. 1C). Several hygromycin-resistant shoots formed after 15 days of culture on R8H65 medium, with a regeneration frequency of 1.6% (Fig. 1D). The hygromycin-resistant shoots were grown well on hormone-free medium (MSRO) and matured into healthy seedlings (Fig. 1E). The seedlings were transferred to soil for acclimatization in the greenhouse (30°C, natural sunlight) (Fig. 1F). To confirm the integration of the T-DNA into the genome, we amplified a piece of transgene sequence and we found that 83.6% of the regenerated plants had successfully integrated the T-DNA sequence into their genomes.

**Fig. 1.**
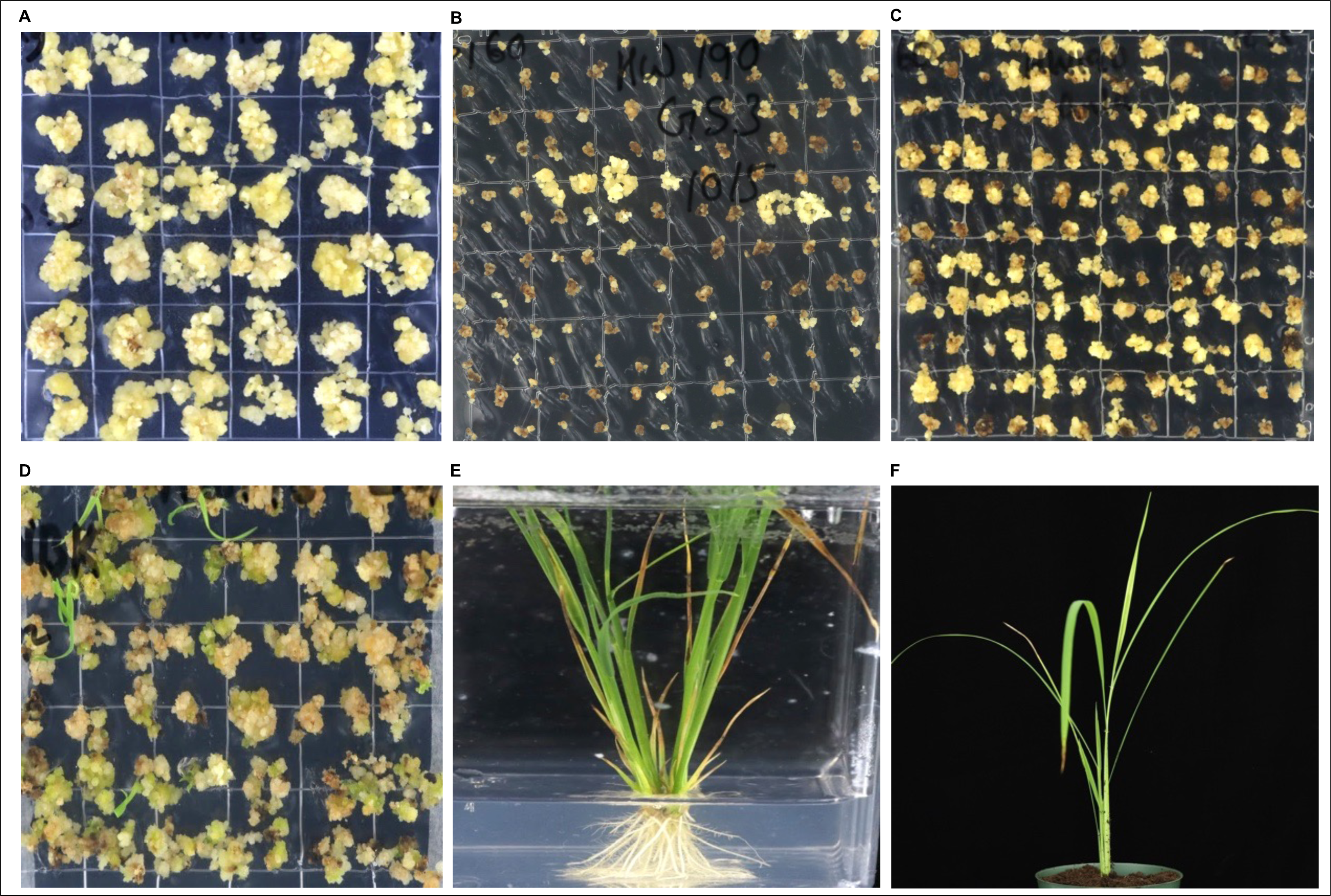
Establishment of *Agrobacterium*-mediated transformation system of *Hassawi* rice. A) Embryogenic-like callus induced from mature grain explants. B) First selection of transformed calli after co-cultivation on NBKCH60 medium containing 50 mg/L hygromycin and 200 mg/L timentin. C) Second selection of the surviving calli on the nNBKCH65 medium containing 65 mg/L hygromycin and 200 mg/L timentin. D) Shoot regeneration from antibiotic-resistant putative transformant calli on R8H65 regeneration medium containing 65 mg/L hygromycin. E) Rooting of putative transformed plantlets on ½ MS and hormonal-free medium containing 65 mg/L hygromycin. F) Soil acclimatization of the plantlets in the greenhouse condition.

### 3.3. Generation of a high-quality reference genome for O. sativa cv. Hassawi

To facilitate trait engineering of Hassawi rice, we attempted to generate the first high-quality reference genome. Therefore, we sequenced the Hassawi genome and generated 539,478 539,478 PacBio HiFi CCS reads (10 Gbp) with an average length of 18.9 Kbp, which corresponds to ∼27x coverage of the total genome. The primary genome assembly consisted of 36 contigs with an N50 = 30.8 Mb (Table 1) and a total size of 391 Mbp. We performed scaffolding using the GPM tool [32] using the *O. sativa* IRGSP Nipponbare RefSeq V1.0 as the reference guide. Results were validated by optical map comparisons, as well dot plot comparisons to the genome of *O. sativa* variety Nipponbare (Supplementary Fig. S2). The final genome assembly was highly contiguous, consisting of 12 pseudomolecules and only nine small gaps (Supplementary Table S3).

**Table 1.**
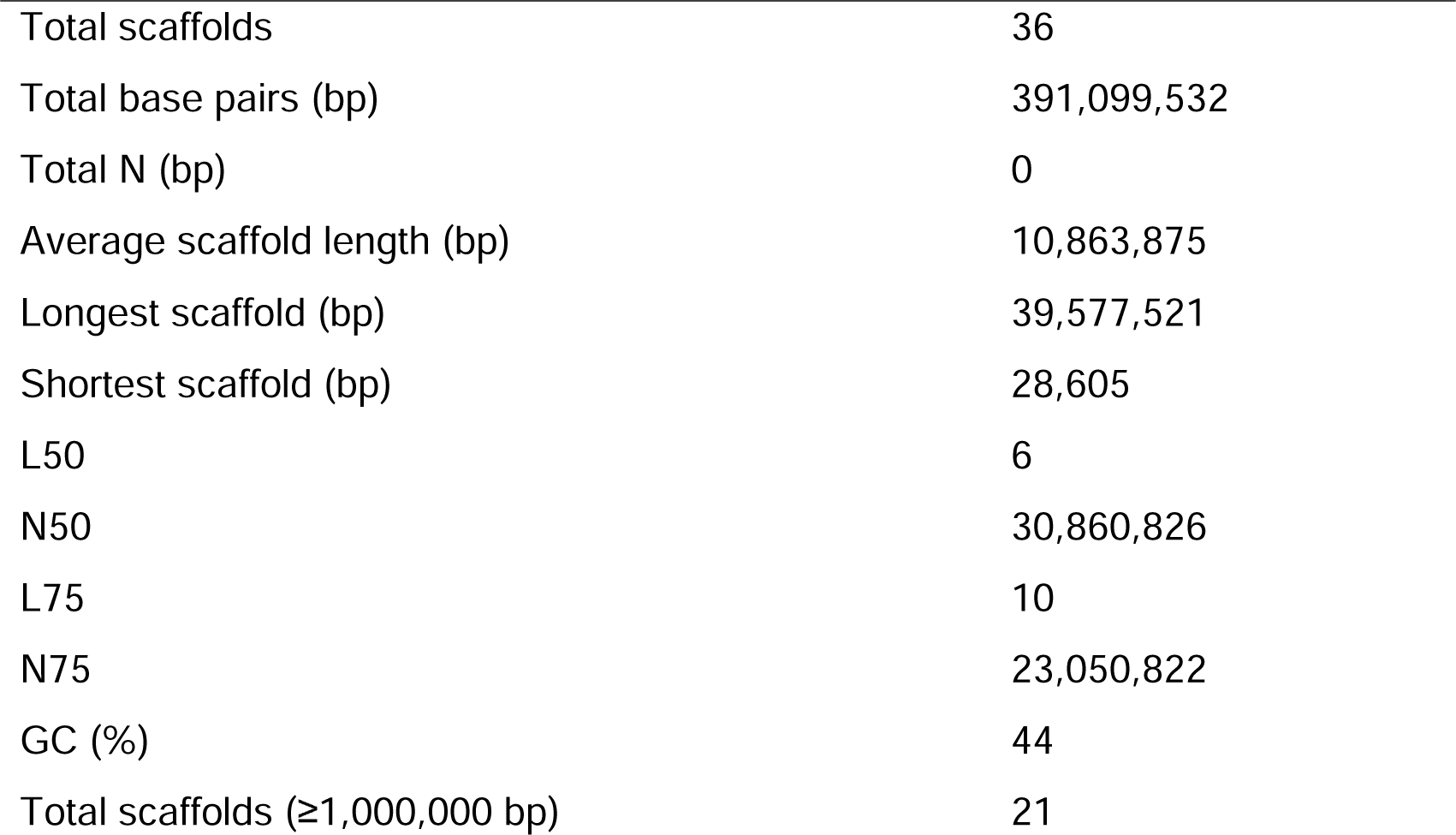
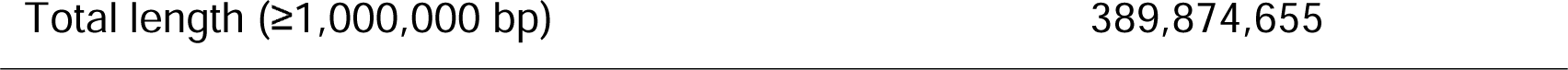
Statistics of the Hassawi genome assembly.

### 3.4. Short-read sequencing of Hassawi (accession 190)

In order to generate the high-quality genome reference sequence of true Hassawi rice, we sequenced the Hassawi 190, the accession used for genome engineering, and carried out the PCA analysis to confirm the genetic groups of Asian rice. Resequencing the Hassawi rice genome using Illumina technology produced 1,681,496,380 reads (151 bp with ∼650 X coverage). We combined these reads with the resequencing data of 160 other accessions, which 10 accessions (>14X) of each of 16 subpopulations from 3K-RGP data set, (Supplementary Table S4) and identified a total of 456,006 SNPs. Principal component analysis (PCA) of these SNPs identified four major subgroups of rice: *circum-Aus* (*cA*), *circum-Basmati* (*cB*), *Geng-japonica* (*GJ*), *Xian-indica* (*XI*) (Fig. 2); Hassawi 190 clustered with *cA*. These results indicate that the Hassawi rice accession examined in this study is from the *cA* subgroup.

**Fig. 2.**
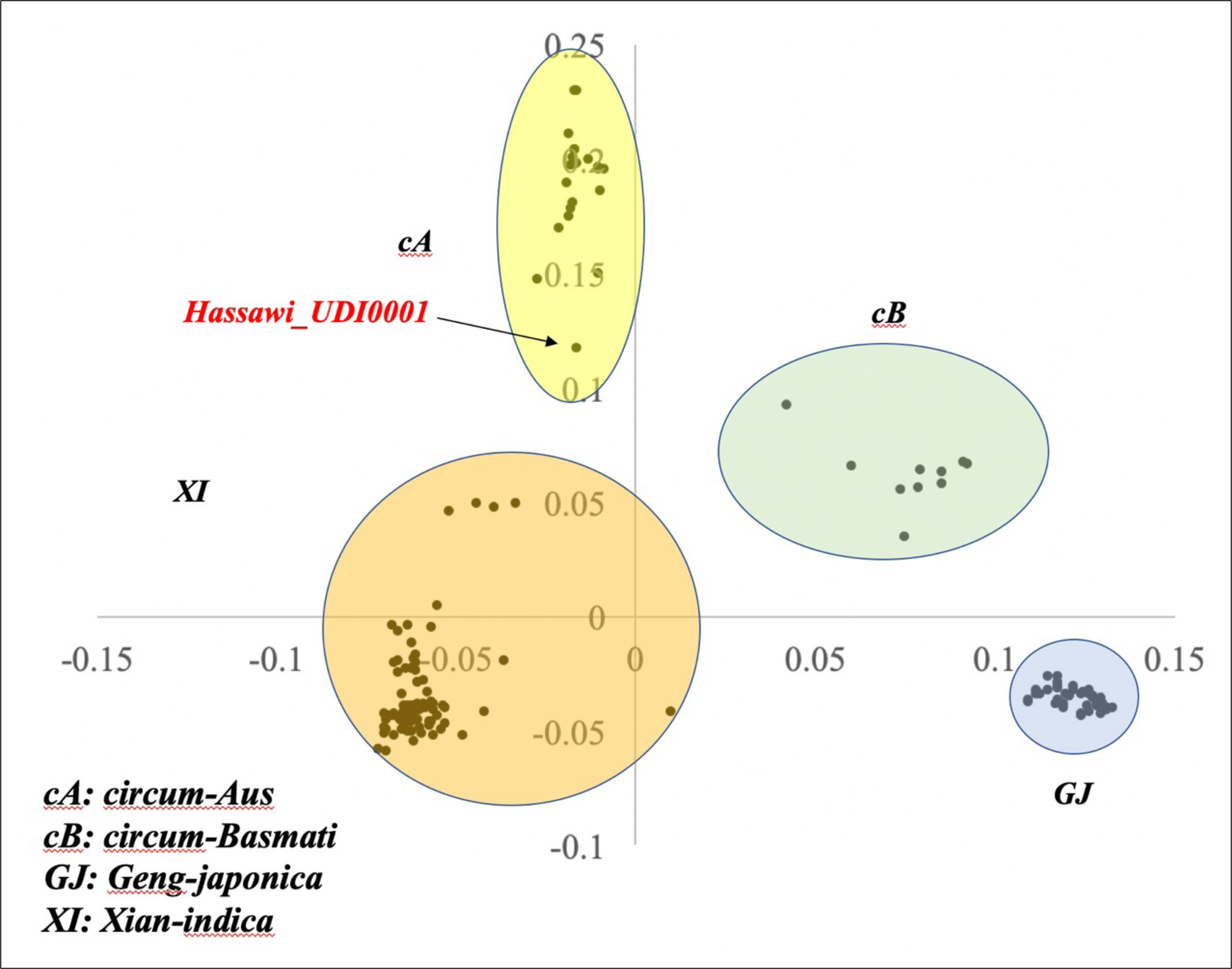
Principal component analysis plot showing the clustering of the Hassawi rice into the four main groups of *O. sativa* (cA: *circum-*Aus, cB: *circum-*Basmati, GJ: *Geng-japonica*, XI: *Xian*-*indica*) The analysis was performed on Hassawi accession 190 and 160 accessions selected from the 3K-RG dataset.

### 3.5. Structural variation (SV) and SNP analysis of the Hassawi genome

We compared the Hassawi genome assembly to the 16 *O. sativa* genomes described in [31] to identify inversions, insertions, and deletions (Supplementary Table S5 and S6). As expected, the number of structural variants (SVs) was larger when Hassawi (which belongs to *O. sativa* subspecies indica) was compared to rice accessions belonging to the subspecies *japonica* vs. *indica*. We used the Hassawi genome assembly to call SNPs and InDels versus the 3K-RGP data [42] and identified 4.3M InDels and 27.6M SNPs. These analyses combined places the Hassawi accession in the *circum-Aus* (*cA*) subpopulation.

### 3.6. Transposable element (TE) analysis

Overall, TEs represent 48.42% of the whole genome assembly, with Ty3-gypsy LTR-RTs (22.24%) and DNA-TEs (18.07%, including MITEs) being the most abundant TE classes. The overall TE content is slightly higher in Hassawi than the rice variety Nipponbare (48.42% vs. 46.65%). Most of the TE classes showed no significant differences in abundance between the two accessions, except for Ty3-gypsy elements, which were slightly more abundant in Hassawi (22.24% vs. 19.23%).

### 3.7. De novo prediction of protein-coding genes

We predicted protein-coding genes in Hassawi using Augustus [55], included within OmicsBox [48], supported by extrinsic Iso-seq and RNA-seq data as evidence. Overall, we identified 36,450 putative genes (Supplementary Table S7) with a BUSCO score of 99.06% (complete single copy: 92%, complete duplicated: 2.2%, fragmented: 4.9%, and missing: 0.94%). The 36,450 *de novo* annotated coding sequences (CDS) were aligned against the 42,355 *O. sativa* IRSPG-1.0 CDSs, which yielded one or multiple hits for 87.1% (31,731 CDSs) of the annotation. These results are similar to those of an *aus* subspecies genome aligned against the Nipponbare reference genome based on our published Rice Gene Index [50]. We detected strong congruence of the *de novo* annotations of Hassawi genes compared to Nipponbare genes across the 12 rice chromosomes (Supplementary Fig. S3).

### 3.8. The CRISPR/Cas9 system knocks out a flowering repressor gene in Hassawi rice at high efficiency

Since we established Hassawi rice regeneration and transformation protocols and generated a high-quality reference genome, we attempted to use CRISPR systems to engineer traits of interest to local farmers including, rapid growth, lodging resistance, high yields, and less water needed for irrigation. To shorten the lifecycle of Hassawi rice, we targeted *Hd4*, a major flowering repressor gene, as functional disruption of this gene was previously shown to accelerate the heading date and reduce the stem length of black rice cv. Cempo Ireng (Sedeek et al 2023). We designed a single guide RNA (sgRNA) to target the second exon of *Hd4* (Fig. 3). We engineered the pRGEB32 vector to express this sgRNA under the control of the *U3* promoter and plant codon-optimized *Cas9* driven by the *Ubiquitin* promoter (Fig. 3A). We transformed the vector into *Agrobacterium tumefaciens* EHA105 and used these cells to transform Hassawi calli using our optimized transformation protocol.

**Fig. 3.**
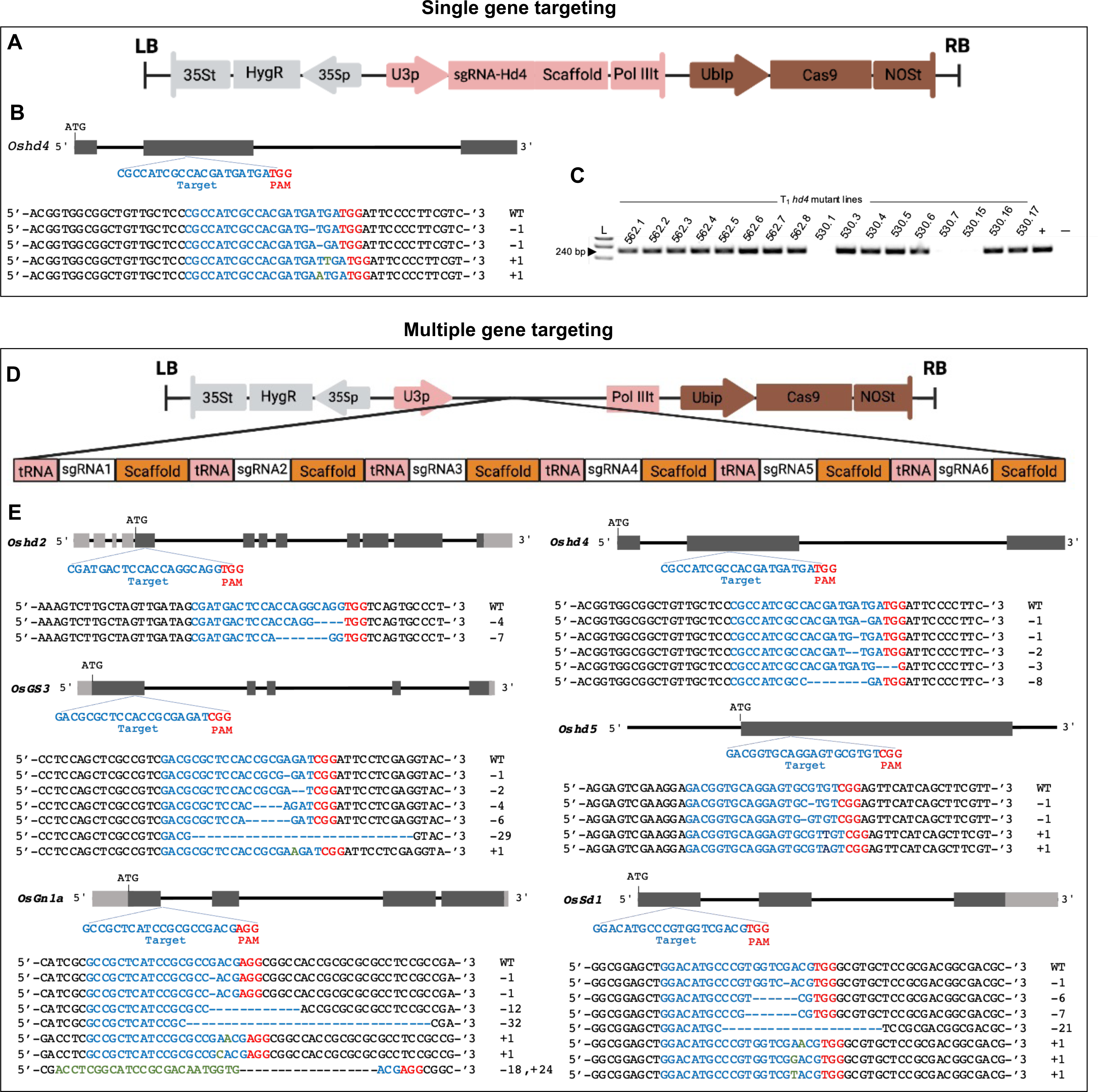
Schematic diagram of single and multiple genetargeting in Hassawi rice. A) Schematic illustrating of the Cas9 and sgRNAs expression cassettes in a single binary vector for rice stable transformation. LB: left border; RB: right border. B) Schematic map of the *hd4* genomic loci in Hassawi rice and site selection for designing sgRNA. The PAM (TGG) is shown in red, sgRNA is shown in red blue. Black boxes indicate exons; black lines indicate introns. A sequence alignment of the target region in the wild-type (WT) and mutant T_2_ lines shows the different insertion/deletions in the mutant lines. C) Gel electrophoresis of the PCR products (240 bp) to select the T-DNA free T_1_ *hd4* engineered Hassawi plants. L; is 1kb plus ladder. D) Schematic illustrating of the T-DNA region containing Cas9 and six sgRNA separated by tRNA in a single binary vector. LB: left border; RB: right border; sgRNA1-6: are for *Hd2, Hd4, GS3, Sd1, Gn1a*, and *Hd5* gene target respectively. E) Schematic map of the *hd2, hd4, hd5, GS3, Gn1a*, and *Sd1* genomic loci in Hassawi rice and site selection for designing sgRNA. The PAM (NGG) is shown in red, sgRNA is shown in red blue. Black boxes indicate exons; black lines indicate introns. A sequence alignment of the target region in the wild-type (WT) and mutant T_2_ lines shows the different insertion/deletions in the mutant lines.

We screened the regenerated T_0_ transformants by PCR to confirm the insertion of the CRISPR materials into the rice genome by specifically amplifying 240 bp of the *Cas9* and Nos terminator sequences (Primer sequences are presented in Supplementary Table S2). A total of 38 independent lines (70% of the regenerated plants) were transgenic. We directly sequenced the targeted flanking regions of 10 randomly selected transgenic plants and detected different types of on-target nucleotide insertions/deletions in eight plants (editing efficiency 80%). Two *hd4* mutants were homozygous for insertions of a single-nucleotide (A) that completely altered the downstream amino acid sequence and created a stop codon that disrupted gene function (Fig. 3B). The remaining mutants were heterozygous (1 line), biallelic (4 lines), or chimeric (1 line) (Supplementary Table S8). To determine whether the desired mutations were transmitted to the next generation and to identify T-DNA-free homozygous mutants, we grew T_0_ and wild-type seeds under greenhouse conditions. We screened the 17 T_1_ progenies by PCR and selected T-DNA-free plants for subsequent genotyping by sequencing. We selected three T-DNA-free homozygous mutant lines for the T_2_ generation (Fig. 3C, and Supplementary Table S8).

### 3.9. hd4 mutants have shortened lifecycles, reduced plant height, and improved yield

We analyzed five homozygous T_2_ *hd4* mutants and five wild-type plants for heading date and other phenotypic characteristics. The panicles of the *hd4* mutants emerged remarkably early (102.2 days of cultivation) compared with the wild-type control (171.6 days) (Fig. 4F). Notably, the plant height of the *hd4* mutants was significantly reduced, measuring only 148.8 cm compared to the wild type (178.8 cm) (Fig.4C, G). Surprisingly, there was a substantial increase (39.1%) in the number of grains per plant in the *hd4* mutants. This could potentially be attributed to the smaller number of grains produced by wild-type plants under greenhouse conditions. Long rice awn present a disadvantage as they possess needle-like structures that cause irritation to farmers’ skin during harvesting and threshing processes. Additionally, several studies have indicated that the loss of awns enhances rice grain number, implying that the presence of rice awns negatively affects overall yield [56, 57]. We observed a significant decrease (of 9.6%) in grain length and a 24.3% reduction in awn length in *hd4* vs. the wild type (Fig. 4D, H). However, no significant differences in tiller number, primary branches per panicle, secondary branches per panicle, spikelets per panicle, grain width, or seed setting rate were detected (Supplementary Fig. S4).

**Fig. 4.**
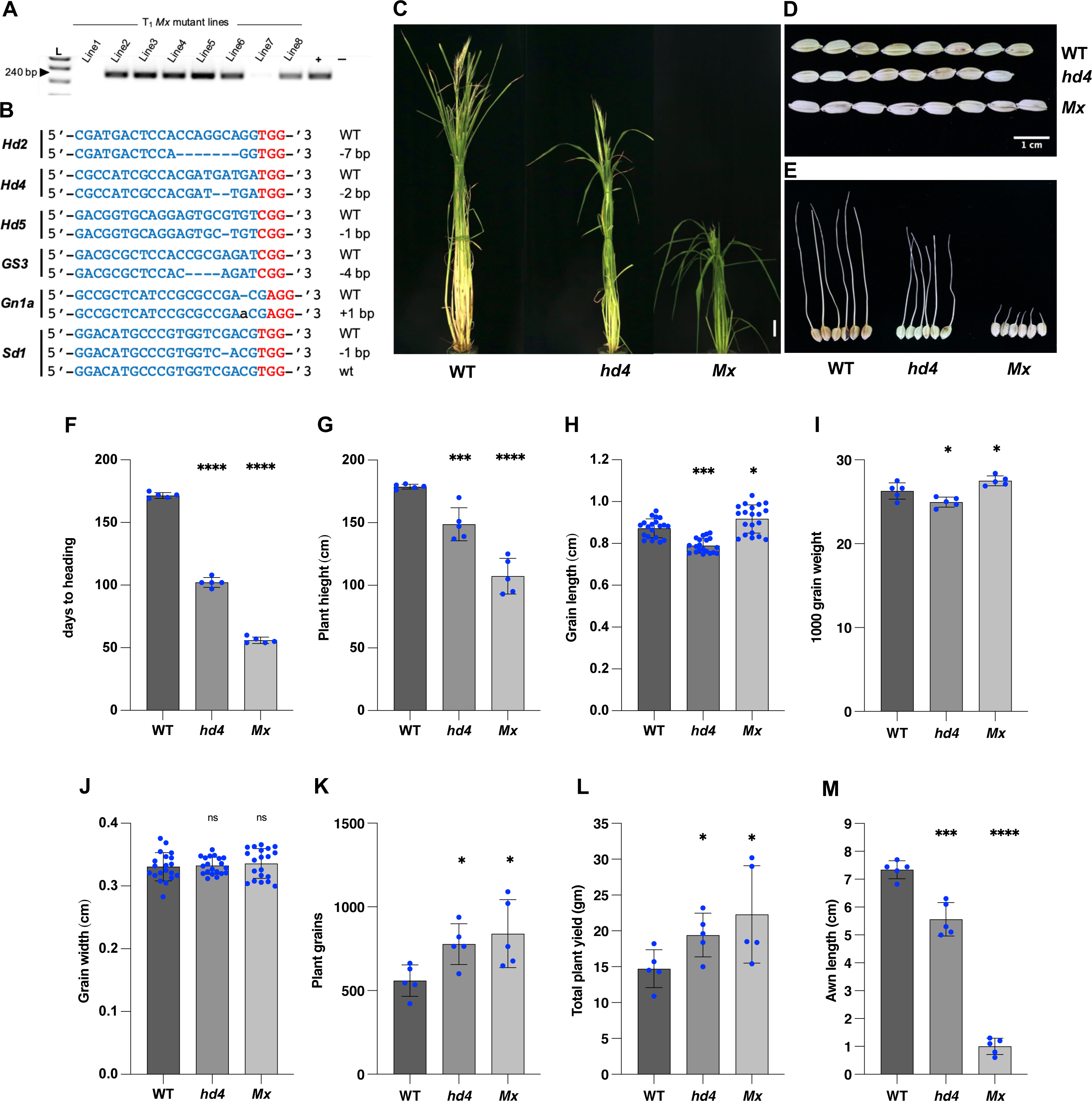
CRISPR/Cas9-induced knockout of Hassawi rice genes and mutant lines phenotyping. A) Gel electrophoresis of the PCR products (240 bp) to select the T-DNA free T_1_ mutants Hassawi plants. L; 1kb ladder. B) Sequence alignment of the target region in the wild-type (WT) and the selected *Mx* T_2_ mutant. C) A representative image of the wild type and *hd4*, *Mx* mutant lines, scale bar, 20 cm. D) Grain phenotypes of the *hd4* and *Mx* mutant lines and WT, scale bar, 1 cm. E) Awn length differences among *hd4* and *Mx* mutant lines and WT. F–M) Unpaired t-test comparing the mutant lines traits and WT (n=5). Data are presented as mean values with the error bars denoting 95% confidence intervals. Asterisks indicate significant differences to the WT: *, p<0.05; ***, p<0.001; ****, p<0.0001. F) Heading date, (n=5). G) Plant height, (n=5). H) Grain length, (n=20). I) Thousand grain weight (n=5). J) Grain width, (n=20). K) Plant grains, (n=5) L) Total number of plant grains (n=5). M) Awn length, (n=5).

### 3.10. Using CRISPR/Cas9 to simultaneously knock out six genes in Hassawi rice

We attempted to simultaneously knock out three major floral repressor genes: *Hd2*, *Hd4*, and *Hd5*. These genes have pleiotropic effects and regulate plant architecture and productivity independently of the flowering pathway. Their functional disruption might impair plant productivity by reducing the number of grains per panicle [58]. Therefore, in addition to these three genes, we targeted *GS3* (a negative regulator of grain size) and *Gn1a* (a negative regulator of grain number) to improve the total productivity of the mutant plants. In addition, we targeted the *Sd1* gene to further reduce plant height and improve lodging resistance. Using our high-quality reference genome, we designed a sgRNA to target the first exon of the coding sequence of *Hd2*, *Hd5*, *GS3, Gn1a*, and *Sd1* and the second exon of *Hd4* (Fig. 3). We cloned all six sgRNAs into a single binary vector (pRGEB32) under the control of a single promoter (*U3*). The sgRNAs were separated by a tRNA sequence that could be digested by endogenous nucleases to release the individual sgRNAs (Fig. 3D). We then transformed embryogenic Hassawi calli with Agrobacterium containing the multiplexed vector (pRGEB32-Mx) and selected transformed cells on medium containing hygromycin.

We obtained 11 hygromycin-resistant plants, four of which were truly transgenic and carried the T-DNA sequence, as confirmed by PCR amplification. Two T_0_ transgenic plants carried different nucleotide insertions/deletions in all six targeted genes that inactivated their functions, but none of the mutant lines were homozygous. The other two plants were quadruple (*hd2/hd4/hd5/gs3*) and pentuple (*hd2/hd4/hd5/gs3/gn1a*) mutants (Supplementary Table S9). We collected the seeds of the mutant lines and grew them and screened eight T_1_ progenies at the seedling stage using PCR and identified two T-DNA plants (Fig. 4A). We sequenced the loci of the six targeted genes but could not recover a homozygous mutant for all six genes. Therefore, we grew the T_1_ seeds and screened eight T_2_ plants by Sanger sequencing (Supplementary Table S9). Only one plant (*Mx*) contained a homozygous mutation in five target genes and a heterozygous mutation in *sd1* (Fig. 4B). We grew the *Mx* seeds for propagation and selected five plants for further phenotypic analysis against the wild-type control.

### 3.11. The Mx mutants developed significantly faster, with enhanced lodging resistance and increased overall yield

We selected five *Mx* and five wild-type plants for phenotypic analysis. The panicles of the *Mx* mutants emerged exceptionally early, i.e., after only 56 days of cultivation, whereas the wild-type plants flowered normally after 171.6 days (Fig. 4F). This remarkably shortened lifecycle was primarily attributed to the simultaneous knockout of three flowering repressors: *Hd2*, *Hd4*, and *Hd5*. Previous studies demonstrated that the loss of function of these repressors suppresses plant elongation in rice. Indeed, we observed a significant reduction in plant height in the *Mx* mutants, measuring 107.4 cm compared to the wild type (178.8 cm) (Fig. 4C, G).

Moreover, the grain length of the *Mx* mutants significantly increased (by 5.1%) due to the knockout of *GS3*, a negative regulator of grain size (Fig. 4D, H). Consequently, the 1000-grain weight of the *Mx* mutants increased significantly (to 27.5 g) compared to wild-type rice (26.3 g) (Fig. 4I). However, no significant difference in grain width was observed between *Mx* and the wild type (Fig. 4J). The overall yield of *Mx* plants exhibited a substantial increase of 51.5% compared to the wild type, primarily driven by a 50.2% increase in grain number compared to the unedited control (Fig. 4K, L). This increased grain number was mainly attributed to the disruption of *Gn1a*, a negative regulator of grain number in rice. Finally, the awn length of *Mx* was significantly reduced (by 86.4%) compared to wild type (Fig. 4E and M). Notably, no significant differences were observed in terms of tiller number, primary branches per panicle, secondary branches per panicle, spikelets per panicle, or seed setting rate (Supplementary Fig. S4).

### 3.12. Untargeted metabolomic profiling of the modified and unmodified Hassawi rice grain

To assess the grain quality post-gene modification, we screened the metabolomic profile of the modified and unmodified rice grains using the Ultrahigh Performance Liquid Chromatography-Tandem Mass Spectroscopy (UPLC-MS/MS). The analysis included three replicates for each line (*hd4*, *Mx*, WT). A total of 465 biomolecules were identified, and 392 matched known chemical structures in the reference library (Fig. 5A). No significant difference was observed in the overall median level among the three lines, with *Mx* being slightly lower and *hd4* being slightly higher than the WT (Fig. 5B). We further investigated all compounds and pathways for significant changes and listed those exceeding a three-fold difference in one or both modified lines relative to WT (Supplementary Table S10). Remarkably, we observed that both modified lines accumulate 11-18 times chlorophyll catabolites, such as pheophytin *a* and pheophorbide in the mature rice grains than WT (Fig. 5C-D). This observation may suggest that the premature onset of flowering reduces the capacity to fully degrade these catabolites in the grain. Additionally, the modified lines displayed a significant elevation in vitamin-related compounds. The higher levels of pantothenate, thiamin, pyridoxamine, delta-tocopherol, and carotene diol derivatives suggest the potential for better nutritional value for the modified lines (Fig. 5E-F, Supplementary Fig. S5). Notably, the modified lines showed 12-15-fold increase in *p*-coumaroyl-serotonin compared to the WT (Fig. 5G). The functional or physiological implication of this compound in plants is unclear, but it has been reported to have significant anti-inflammatory and anti-cancer properties in human cell studies [59, 60], and other positive health-related effects in experimental animal systems [61].

**Fig. 5.**
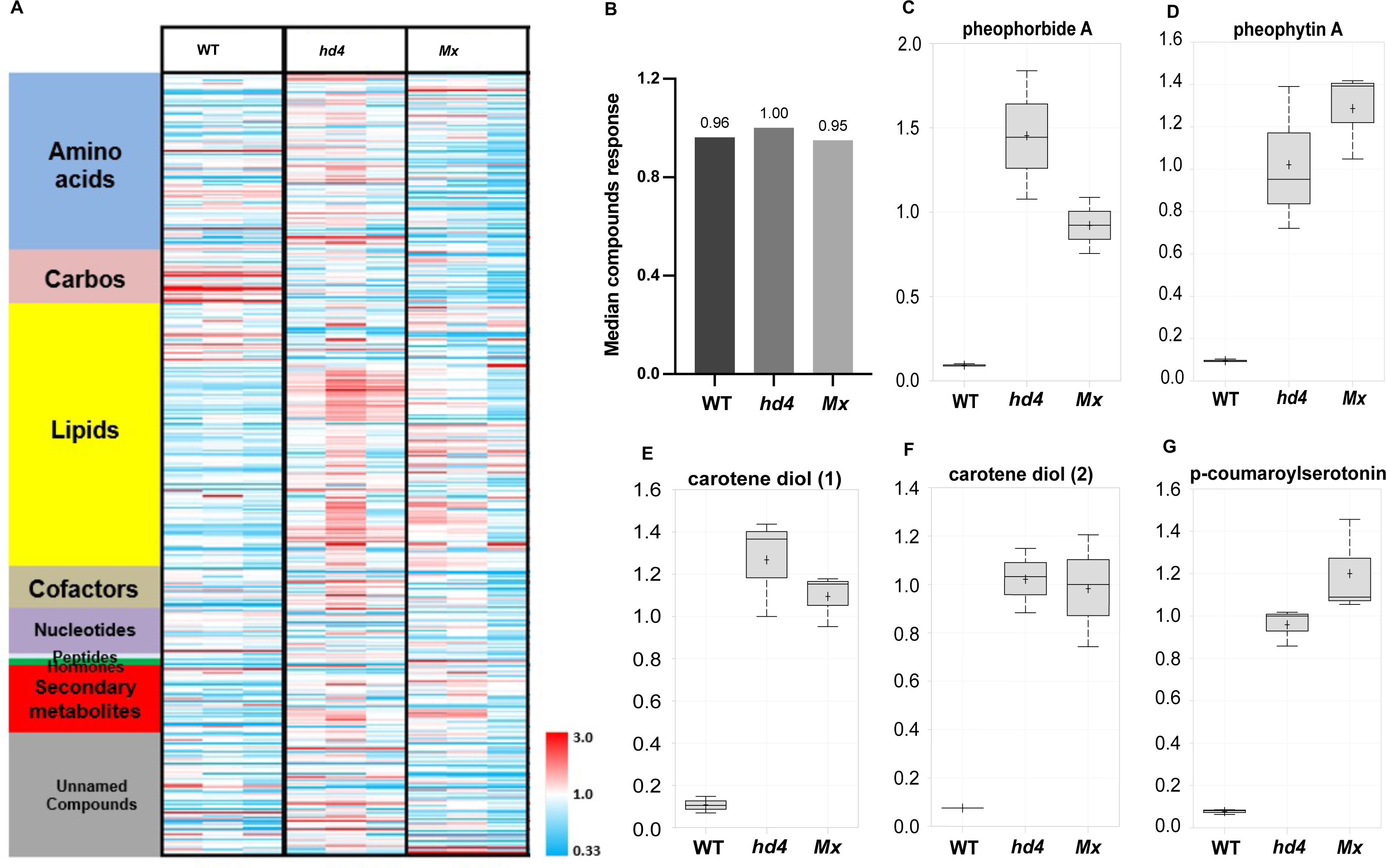
Metabolomic profiling of the modified and unmodified Hassawi rice. A) Heat map visualization of the median scaled imputed data for red rice grain. Red cells indicate levels higher than the median value, while blue cells indicate levels below the median. White cells are near the median. B) Overall median response for all named compounds in red rice, by experimental group. C-G) Box plots of some compounds that highly and significantly different between WT and one or both modified lines.

Significant changes were also observed in the major class of hydroxycinnamic acid esters, the chlorogenic acids (Supplementary Fig. S5). Chlorogenic acids are widely reported to have beneficial effects for cardiovascular, liver, and kidney health [62], as well as to promote cognitive and neuroprotection [63]. We did not detect chlorogenic acid itself in this study, but instead measured a variety of isomers, of these feruloylquinate and *p*-coumaroylquinate esters were significantly higher in one or both of the modified lines compared to WT (Supplementary Fig. S5). Red rice is recognized for its richness in flavonoid compounds, many of which were detected in our study have been widely studied for their antioxidant, anti-inflammatory, and cardiovascular health [64]. However, while several flavonoids (catechin, naringenin, dihydrokaempferol, and quercetin 3-galactoside) were significantly higher than WT in one or both modified lines (Supplementary Fig. S5), only catechin showed a large increase in relation to the gene modifications, being about 3-5 fold higher than WT. Salicylate and its glucoside form, known for its role in plant disease protection and human heart health, were both significantly higher in the modified lines than in the WT (Supplementary Fig. S5).

## 4. Discussion

Hassawi rice grains are nutrient-dense, as they contain a wealth of bioactive compounds and essential nutrients crucial for maintaining good health. The cultivation of Hassawi rice has the potential to enhance food security and mitigate the risks of malnutrition. However, its high cost renders it unaffordable for most consumers, primarily due to its limited cultivation in the water limited region of Al Ahsa. Several factors hinder the widespread cultivation of Hassawi rice: (1) farmers have little interest in growing this rice due to its lengthy growth cycle, low yield capacity, and suboptimal plant height; (2) water resources in Saudi Arabia are severely constrained, making it challenging to allocate enough water for rice cultivation; and (3) the government has restricted the expansion of rice cultivation, as it requires a huge amount of water. One viable solution to addressing these challenges involves shortening the lifecycle of Hassawi rice. By shortening its growth duration, water usage for irrigation can be minimized, potentially encouraging farmers and the government to engage in doubling or tripling of rice cultivation for the same amount of water. The CRISPR/Cas9 system offers a practical, straightforward approach for simultaneously editing multiple targets, enabling the modification of various undesirable traits within the same plant [52, 65].

The precision gene editing and engineering of Hassawi rice requires a full genome sequence and established regeneration and transformation methods to enable the delivery of the CRISPR machinery into plant cells for targeted genome modification. In the current study, we sequenced, assembled, and provided the first comprehensive reference genome of Hassawi rice and assigned it to a specific rice subpopulation (i.e. Aus). These genomic data are essential for future efforts to improve this valuable rice accession. Although efficient regeneration and transformation techniques have been established for the *O. sativa japonica/indica* varietal groups, these methods are genotype-dependent. Nevertheless, optimizing these processes for Hassawi rice holds the key to unlocking its genome engineering potential to improve traits of agronomic and nutritional importance by breeding. Regeneration is the most crucial step in plant transformation via somatic embryogenesis. Therefore, the ability to induce callus and to regenerate shoots and roots from callus is essential for establishing a robust genetic transformation system.

Here we established regeneration and transformation systems for Hassawi rice to enable the targeted genome engineering of traits of value, such as accelerating the heading date, reducing plant height, and improving the overall yield. We individually knocked out one of the main repressors of rice heading date (*Hd4*). The resulting mutants were early maturing and shorter than the wild type. The loss of function of *hd4* was previously shown to negatively affect the number of grains per panicle. Surprisingly, the number of grains per panicle increased in the Hassawi rice *hd4* mutants compared to the wild type. This could potentially be attributed to the smaller number of grains produced by wild-type plants under greenhouse conditions. Therefore, the *hd4* mutant lines must be evaluated in the field under natural cultivation conditions compared to the wild-type control. We successfully knocked out six genes simultaneously to modify many traits of interest in Hassawi rice. The simultaneous targeting of six genes resulted in extremely early maturing, short, high-yielding plants. The new modified lines, especially *Mx*, have many desirable traits that will certainly encourage farmers/investors to cultivate Hassawi on a large scale and generate more profit. The modified lines also had higher amounts of several vitamin-related compounds, as well as increased levels of various secondary metabolites, including flavonoids and chlorogenic acids.

## 5. Conclusion

The nutritional richness of Hassawi rice presents promise for enhancing food security and countering malnutrition. However, challenges like high cultivation costs, water scarcity, and governmental constraints limit its widespread growth. Addressing these issues involves shortening the growth cycle to conserve water and encourage expanded cultivation. Leveraging CRISPR/Cas9 technology, we targeted a set of key rice genes to accelerate maturity and enhance yield. Notably, mutants displayed improved traits, suggesting their potential for large-scale cultivation. Our findings represent a major step towards establishing Hassawi rice as a superfood, which will improve human health and reduce the burden of malnutrition in developing and developed countries.

## Data availability

The PacBio raw data and genome sequences were deposited at NCBI under BioProject ID: PRJNA657951.

## Acknowledgments

We thank the King Abdullah University of Science and Technology for providing the Hassawi rice grains. We thank Haroon Butt, Norhan Hassan, and all members of the Laboratory for Genome Engineering and Synthetic Biology at KAUST for critical discussion and technical help with this work.

## Funding

This work is funded by KAUST-baseline funding to Magdy Mahfouz and Rod Wing.

## Competing interests

The authors declare that they have no competing interests.

## Author contributions

MM, KS, NM, and RW conceived the project; KS, NM, MT, KrS, and SK conducted the experiments; KS, NM, YZ, AZ, and NA analyzed the data, and KS, NM, MM, AZ, YZ, and RW wrote the manuscript.

## Notes

### Competing Interest Statement

The authors have declared no competing interest.

