## Supplementary Information for "Multitrait engineering of Hassawi red rice for sustainable cultivation"

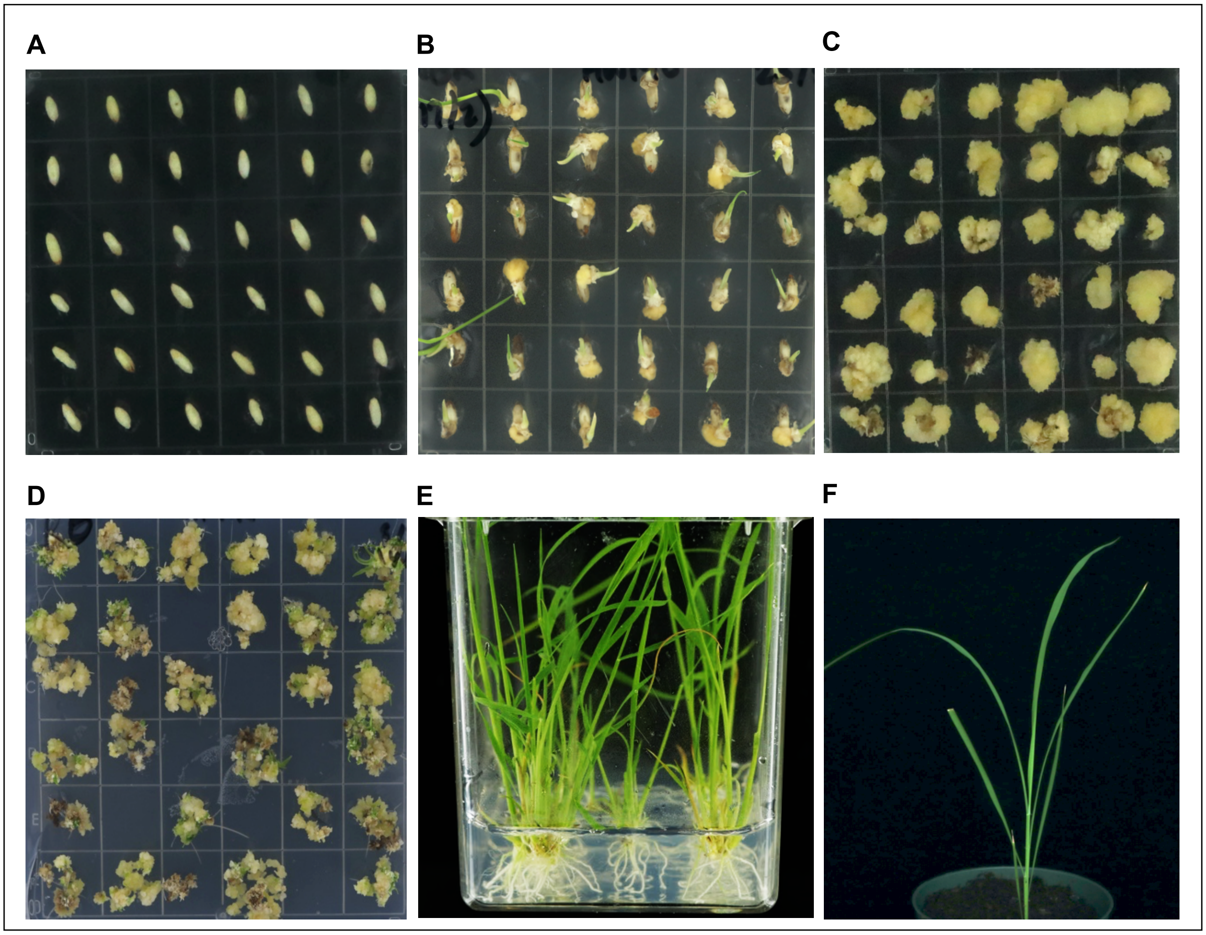


**Supplementary Fig. S1. Main steps of the establishment of a regeneration protocol for Hassawi red rice.** A) Rice mature grains (an explant for callus induction) on callus induction medium. B) Scutella tissue emerged from the grains after 7 days on 2NBK medium. C) Callus propagation on fresh 2NBK medium. D) Shoot induction using R8 medium. E) Rooting on MSRO medium. F) Soil acclimatization of the regenerated plants under the greenhouse condition.

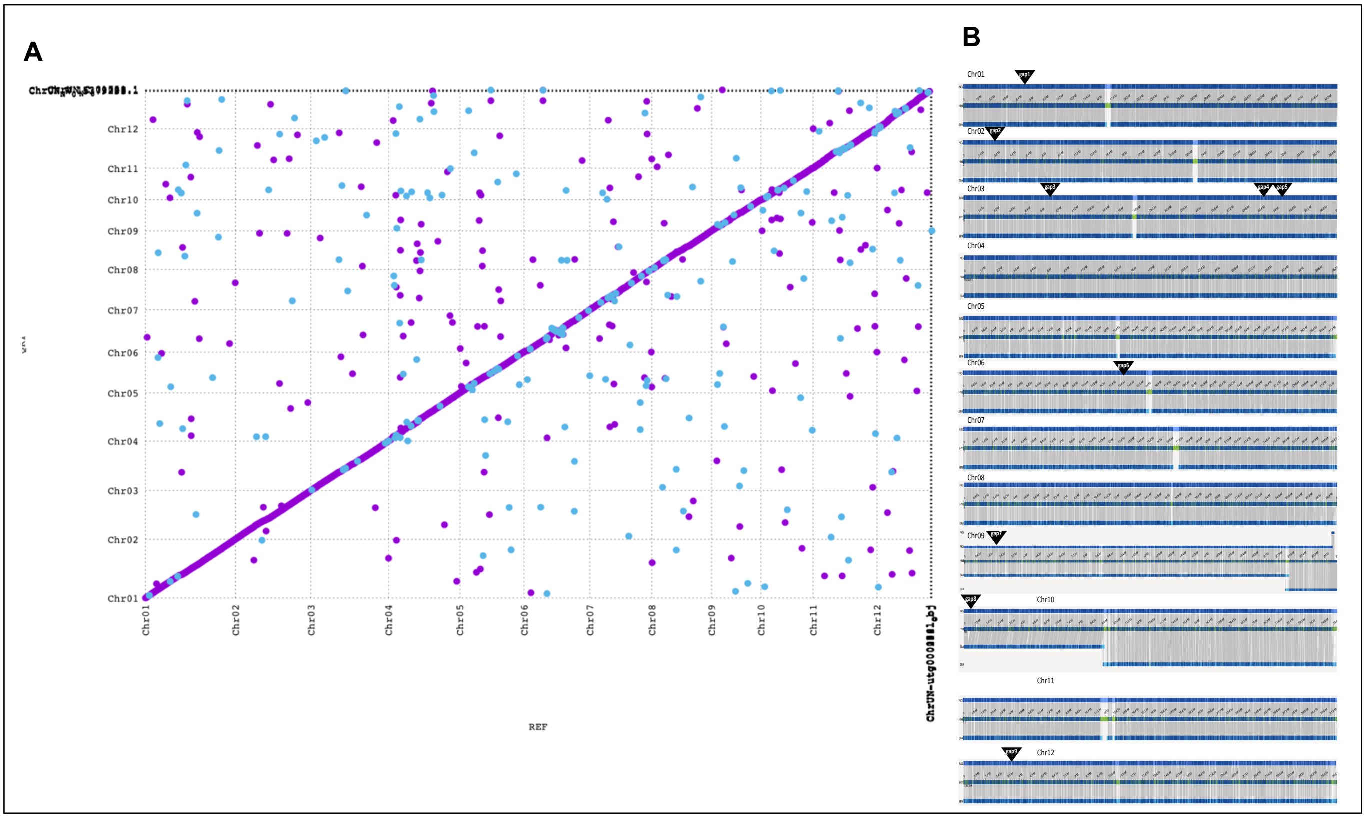

**Supplementary Fig. S2. Validation of the Hassawi genome sequence**. A) dot plot that compares the Hassawi genome to Nipponbare, B) optical map.

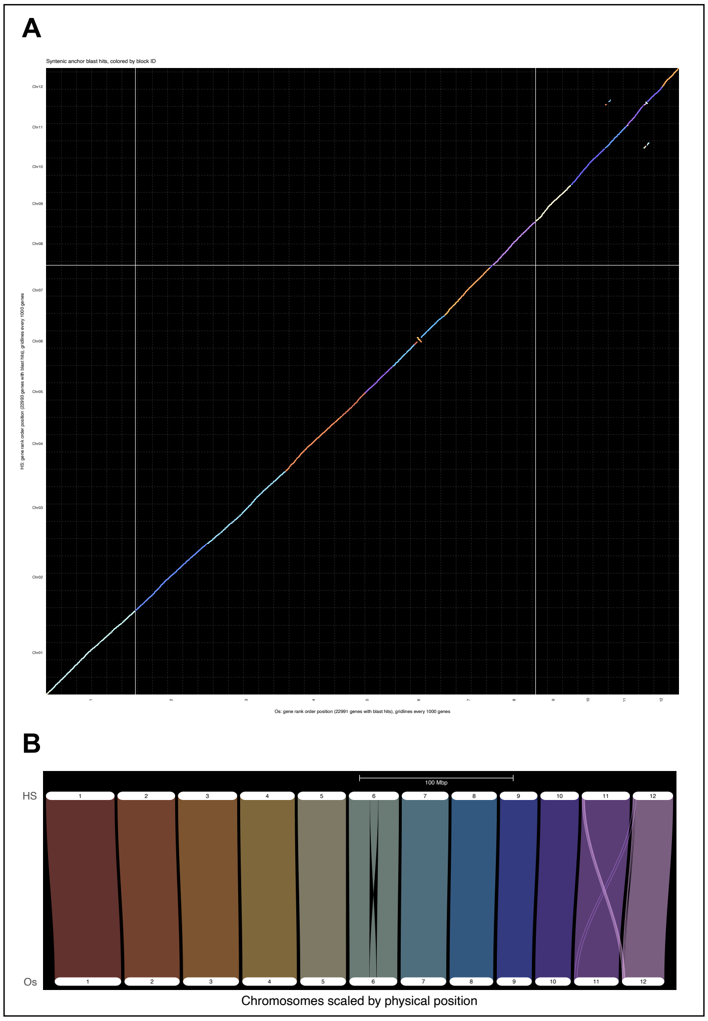


**Supplementary Fig. S3. Synteny and homology analyses reveal a conservation of 87% of Hassawi protein-coding regions against Nipponbare protein-coding regions.** A) The y-axis denotes Hassawi and the x-axis indicates Nipponbare chromosomes. Each dot represents a putative syntenically conserved gene, each color represents a chromosome. B) Syntenic view of Hassawi protein-coding genes across the 12 Nipponbare chromosomes. HS represents Hassawi, and Os represents *Oryza sativa* cv. Nipponbare.

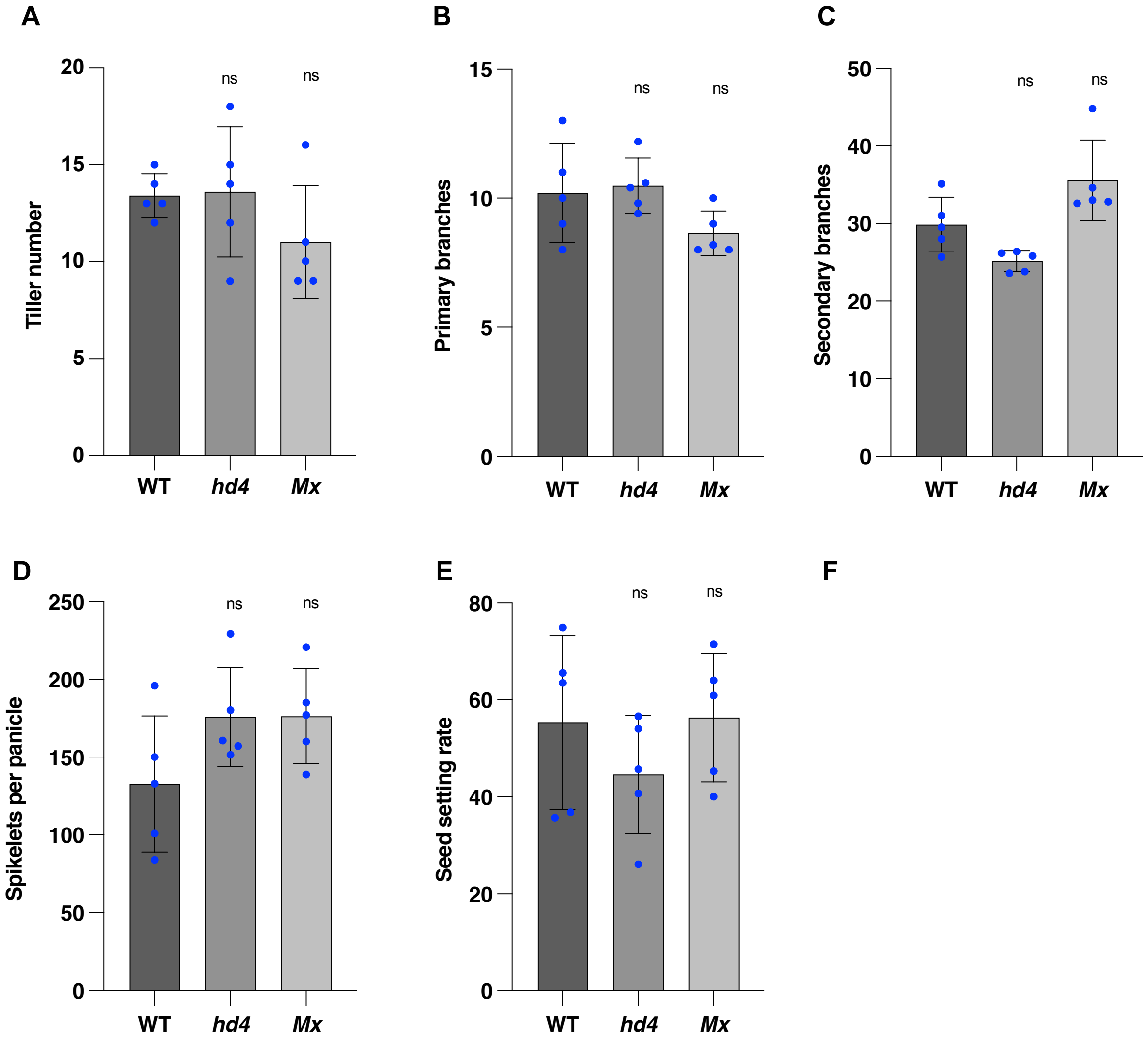


**Supplementary Fig. S4. Phenotyping of the *hd4* and *Mx* mutant lines.** A-F) Unpaired t-test comparing the different yield-related traits of the mutant lines. Data are presented as mean values, with the error bars denoting 95% confidence intervals. Asterisks indicate significant differences from the WT: ns, statistically non-significant. A) Tiller number (n=5). B) Primary branches per panicle (n=5). C) Secondary branches per panicle (n=5). D) Spikelets per panicle, (n=5). E) Seed setting rate (n=5).

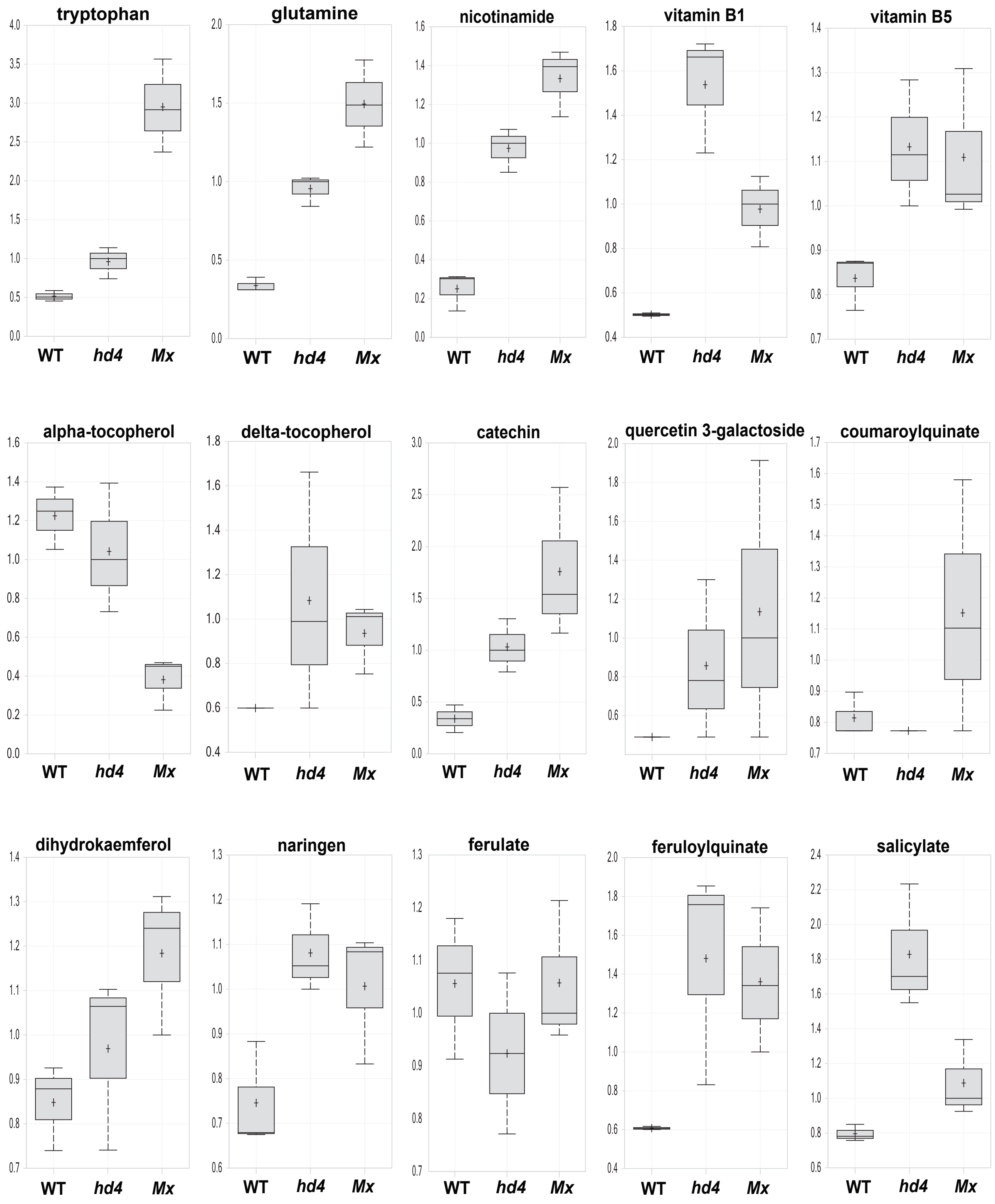


**Supplementary Fig. S5.** Box plots for selected compounds in red rice.

**Supplementary Table S1: Tissue culture media used in this study for callus induction, selection, regeneration, and rooting**

| Medium | Components |
|---|---|
| **2NBK** | 100 mL of 10× N6K major salts + 10 mL of 100× B5 minor salts + 10 mL of 100× B5 vitamins + 10 mL of 100× FeEDTA + 20 mL of 100 mg 2,4-D + 0.5 g L-proline + 0.5 g casein hydrolysate + 30 g maltose + $H_2O$ to 900 mL + pH to 5.8 + 4 g phytagel + 100 mL of 10× AA amino acids (pH 5.8) (add after autoclaving and cooling down to 60°C) |
| **NBKCH50** | 2NBK + 1 mL of 50 mg/mL hygromycin B + 1 mL of 200 mg/mL timentin |
| **nNBKC** | 100 mL of 10× N6K major salts + 10 mL of 100× B5 minor salts + 10 mL of 100× B5 vitamins + 10 mL of 100× FeEDTA + 10 mL of 100 mg 2,4-D + 0.5 mg 1-naphthaleneacetic acid (NAA) + and 0.1 mg 6-benzylaminopurine (BAP) + 0.5 g L-proline + 0.5 g casein hydrolysate + 20 g sucrose + 55 gm D-sorbitol + $H_2O$ to 1000 mL + pH to 5.8 + 5 g phytagel + autoclave at 121°C for 20 min. |
| **nNBKCH65** | nNBKC + 1.3 mL of 50 mg/mL hygromycin B + 1 mL of 200 mg/mL timentin |
| **R8** | 4.3 g MS basal medium + 30 g sucrose + 2 mg BAP + 1 mg NAA + $H_2O$ to 1000 mL + pH to 5.8 + 4 g phytagel + autoclave at 121°C for 20 min. |
| **R8H65** | R8 + 1.3 mL of 50 mg/mL hygromycin B + 1 mL of 200 mg/mL timentin |
| **MSRO** | 2.2 g MS basal medium + 10 g sucrose + 0.5 g casein hydrolysate + $H_2O$ to 1000 mL + pH to 5.8 + 4 g phytagel + autoclave at 121°C for 20 min. |
| **MSROH65** | MSRO + 1.3 mL of 50 mg/mL hygromycin B |

**Supplementary Table S2. List of oligos.** Table lists the oligo sequences used for designing the sgRNAs for the targeted genes and the primers used for detecting the T-DNA insertion and mutant genotypes. Blue nucleotides are the BsaI restriction overhang.

| Name | Sequence | Usage |
|---|---|---|
| Hd2_sgRNA1_F | GGCACGATGACTCCACCAGGCAGG | Design Hd2 sgRNA |
| Hd2_sgRNA1_R | AAACCCTGCCTGGTGGAGTCATCG | Design Hd2 sgRNA |
| Hd4_sgRNA1_F | GGCACGCCATCGCCACGATGATGA | Design Hd4 sgRNA |
| Hd4_sgRNA1_R | AAACTCATCATCGTGGCGATGGCG | Design Hd4 sgRNA |
| Hd5_sgRNA1_F | GGCAGACGGTGCAGGAGTGCGTGT | Design Hd5 sgRNA |
| Hd5_sgRNA1_R | AAACACACGCACTCCTGCACCGTC | Design Hd5 sgRNA |
| GS3_sgRNA1_F | GGCAGACGCGCTCCACCGCGAGAT | Design GS3sgRNA |
| GS3_sgRNA1_R | AAACATCTCGCGGTGGAGCGCGTC | Design GS3 sgRNA |
| Gn1a_sgRNA1_F | GGCAGCCGCTCATCCGCGCCGACG | Design Gn1a sgRNA |
| Gn1a_sgRNA1_R | AAACCGTCGGCGCGGATGAGCGGC | Design Gn1a sgRNA |
| Sd1_sgRNA1_F | GGCAGGACATGCCCGTGGTCGACG | Design Sd1 sgRNA |
| Sd1_sgRNA1_R | AAACCGTCGACCACGGGCATGTCC | Design Sd1 sgRNA |
| Hd2_F1 | ATGATGGGAACCGCTCATCAC | Amplify Hd2 target region |
| Hd2_R6 | CATACATGCAGTGACGAAGC | Amplify Hd2 target region |
| Hd4_F2 | ATGTCGATGGGACCAGCAGC | Amplify Hd4 target region |
| Hd4_R3 | GATGGTGGCGCTGGCCGCGG | Amplify Hd4 target region |
| Hd5_F2 | ACTTGCTGAGCCCGGTGGGC | Amplify Hd5 target region |
| Hd5_R3 | GCGTGGTCATGGCCCAGAGG | Amplify Hd5 target region |
| GS3_UF2 | TATCGGAACTTCGGAGTGAC | Amplify GS3 target region |
| GS3_R8 | TGATGCATGCATGCAGAAGC | Amplify GS3 target region |
| Gn1a_F15 | TTCGTCCGCCTCCTTCCTCG | Amplify Gn1a target region |
| Gn1a_R17 | TGCACCGAGTGGCCACACCC | Amplify Gn1a target region |
| Sd1_F2 | ACGCTCTCAACTCACTCCCGC | Amplify Sd1 target region |
| Sd1_R3 | ACCTGGAAGAACCCGTGCGTG | Amplify Sd1 target region |
| Cas9_F7 | GATCGACCTGTCTCAGCTGG | Amplify 240 bp of the T-DNA sequence |
| Nos R7 | CGGCAACAGGATTCAATCTTAAG | Amplify 240 bp of the T-DNA sequence |

**Supplementary Table S3. The nine gaps observed in the Hassawi reference genome.**

| Chromosome number | From | To | Gap name |
|---|---|---|---|
| Chr01 | 5,111,280 | 5,111,379 | Name=gap1; size=100 |
| Chr02 | 1,919,222 | 1,919,321 | Name=gap2; size=100 |
| Chr03 | 8,723,576 | 8,723,675 | Name=gap3; size=100 |
| Chr03 | 30,641,432 | 30,641,531 | Name=gap4; size=100 |
| Chr03 | 32,172,588 | 32,172,687 | Name=gap5; size=100 |
| Chr06 | 12,802,555 | 12,802,654 | Name=gap6; size=100 |
| Chr09 | 23,050,823 | 23,050,922 | Name=gap7; size=100 |
| Chr10 | 1,641,307 | 1,641,406 | Name=gap8; size=100 |
| Chr12 | 3,386,367 | 3,386,466 | Name=gap9; size=100 |

**Supplementary Table S4. PCA of 161 rice accessions used to confirm the identity of Hassawi rice used in transformation.**

| Accession | NCBI | PCA1 | PCA2 | PCA3 | Subpopulation |
|---|---|---|---|---|---|
| IRIS_313-11513 | ERS468722 | -0.0714834 | -0.0582574 | 0.00429076 | XI-1B2 |
| IRIS_313-8616 | ERS467956 | -0.0700572 | -0.0423573 | 0.00391561 | XI-1B1 |
| CX230 | ERS470554 | -0.0699772 | -0.0491165 | 0.00585043 | XI-1B1 |
| IRIS_313-11941 | ERS469086 | -0.0697796 | -0.0513258 | 0.00425711 | XI-1B2 |
| IRIS_313-7807 | ERS468569 | -0.0695393 | -0.0434523 | 0.00506934 | XI-1B1 |
| IRIS_313-12234 | ERS469271 | -0.0693083 | -0.0593385 | 0.00440728 | XI-1B2 |
| IRIS_313-7816 | ERS468567 | -0.0692704 | -0.0451132 | 0.00617309 | XI-1B1 |
| CX403 | ERS470689 | -0.0692115 | -0.0493634 | 0.00725777 | XI-1B1 |
| IRIS_313-10915 | ERS469713 | -0.0688639 | -0.0435688 | 0.00918521 | XI-3B1 |
| IRIS_313-8660 | ERS467837 | -0.0677618 | -0.00383862 | -0.0111129 | XI-2B |
| IRIS_313-11708 | ERS468872 | -0.0677423 | -0.0432034 | 0.00594083 | XI-3B1 |
| IRIS_313-11072 | ERS469874 | -0.0674495 | -0.043019 | 0.00563662 | XI-3B1 |
| CX83 | ERS470736 | -0.0670357 | -0.052155 | 0.00538577 | XI-1B1 |
| IRIS_313-11418 | ERS470209 | -0.067005 | -0.020199 | 0.00721692 | XI-2A |
| IRIS_313-11841 | ERS468994 | -0.0665289 | -0.0439275 | 0.00335876 | XI-3B1 |
| IRIS_313-11358 | ERS470152 | -0.0663298 | -0.0250638 | 0.00456802 | XI-2A |
| IRIS_313-11638 | ERS468814 | -0.0659709 | -0.00647023 | -0.00309306 | XI-2B |
| IRIS_313-11370 | ERS470163 | -0.065939 | -0.0197203 | -0.000523082 | XI-2B |
| IRIS_313-11512 | ERS468721 | -0.0656909 | -0.0451581 | 0.00464713 | XI-3B2 |
| IRIS_313-10924 | ERS469723 | -0.0652243 | -0.0337342 | 0.00257388 | XI-adm |
| B104 | ERS470314 | -0.0651814 | -0.0493329 | 0.000145058 | XI-1A |
| IRIS_313-11860 | ERS469013 | -0.0648586 | -0.0440927 | 0.0016107 | XI-1A |
| IRIS_313-9005 | ERS468273 | -0.06477 | -0.0424697 | 0.00878977 | XI-3A |
| IRIS_313-11921 | ERS469065 | -0.0647206 | -0.0419847 | 0.00826615 | XI-3B1 |
| IRIS_313-9372 | ERS468063 | -0.0646318 | -0.0391669 | -0.00149144 | XI-1A |
| CX53 | ERS470703 | -0.0646274 | -0.0475732 | 0.00185955 | XI-adm |
| IRIS_313-10791 | ERS469600 | -0.0646076 | -0.049857 | 0.0048242 | XI-3A |
| IRIS_313-10997 | ERS469793 | -0.0643148 | -0.0470927 | 0.00252255 | XI-3A |
| IRIS_313-11369 | ERS470162 | -0.0637832 | -0.0232619 | 0.0113514 | XI-2A |
| CX76 | ERS470729 | -0.0636847 | -0.045449 | 0.00169525 | XI-1B1 |
| CX26 | ERS470574 | -0.0634382 | -0.0396436 | 0.00573869 | XI-1B1 |
| IRIS_313-8466 | ERS468244 | -0.0634347 | -0.0448811 | 0.00500909 | XI-3A |
| CX421 | ERS804456 | -0.0633404 | -0.0503478 | 0.0024197 | XI-1B2 |
| IRIS_313-11895 | ERS469037 | -0.0632887 | -0.0411859 | 0.00912031 | XI-3B1 |

| | | | | | |
|---|---|---|---|---|---|
| IRIS_313-10721 | ERS469527 | -0.0632726 | -0.00410929 | -0.012642 | XI-2B |
| IRIS_313-10234 | ERS468178 | -0.0630947 | -0.0465321 | 0.00484879 | XI-1B1 |
| CX418 | ERS804453 | -0.0628587 | -0.0427592 | -0.00149101 | XI-1B2 |
| IRIS_313-10970 | ERS469772 | -0.0628229 | -0.0429212 | 0.00945648 | XI-3B1 |
| IRIS_313-10221 | ERS468176 | -0.0627142 | -0.0447133 | 0.00629062 | XI-3B2 |
| IRIS_313-10835 | ERS469638 | -0.0626077 | -0.0113745 | -0.00525725 | XI-2B |
| IRIS_313-11850 | ERS469002 | -0.0625013 | -0.0433835 | 0.0118036 | XI-3A |
| CX424 | ERS804459 | -0.0624867 | -0.0454573 | 0.000866321 | XI-1B2 |
| IRIS_313-11231 | ERS470023 | -0.0624406 | -0.0233019 | 0.0134162 | XI-2A |
| B114 | ERS470324 | -0.0624396 | -0.0500733 | -0.000291871 | XI-1A |
| IRIS_313-11780 | ERS468937 | -0.0621928 | -0.0392267 | 0.00360261 | XI-3B1 |
| B232 | ERS470431 | -0.0621648 | -0.0463409 | 0.00355942 | XI-1B2 |
| IRIS_313-10756 | ERS469565 | -0.0620679 | -0.0188686 | 0.0105996 | XI-2A |
| IRIS_313-11480 | ERS468691 | -0.0620553 | -0.0549965 | 0.00172757 | XI-1B2 |
| IRIS_313-10650 | ERS469470 | -0.061959 | -0.041691 | 0.0103428 | XI-3A |
| IRIS_313-8922 | ERS468271 | -0.0619206 | -0.0225034 | 0.0128884 | XI-2A |
| IRIS_313-9758 | ERS468113 | -0.0617824 | -0.0422468 | 0.00201796 | XI-1A |
| IRIS_313-12273 | ERS469292 | -0.0615829 | -0.0393511 | 0.00316804 | XI-1A |
| IRIS_313-11225 | ERS470017 | -0.0613345 | -0.0238706 | 0.0152598 | XI-2A |
| IRIS_313-11310 | ERS470101 | -0.061326 | -0.0166073 | -0.00162724 | XI-adm |
| IRIS_313-10985 | ERS469789 | -0.0612987 | -0.0200066 | 0.00713862 | XI-2A |
| IRIS_313-12048 | ERS469180 | -0.0612155 | -0.0396811 | 0.0107656 | XI-3A |
| IRIS_313-11952 | ERS469097 | -0.0611922 | -0.0409767 | -0.00110239 | XI-1A |
| IRIS_313-12249 | ERS469280 | -0.0606821 | -0.0291901 | 0.0012215 | XI-3B1 |
| IRIS_313-12296 | ERS469311 | -0.0606364 | -0.0382444 | 0.00501869 | XI-adm |
| IRIS_313-11695 | ERS468870 | -0.0606117 | -0.0456321 | 0.00139796 | XI-adm |
| CX267 | ERS470579 | -0.0604604 | -0.0286924 | -0.00164354 | XI-adm |
| IRIS_313-11182 | ERS469977 | -0.0603898 | -0.048638 | 0.00458239 | XI-3A |
| IRIS_313-12275 | ERS469293 | -0.0600533 | -0.0409952 | 0.00549984 | XI-3B2 |
| CX369 | ERS470654 | -0.0595517 | -0.0512507 | 0.00215255 | XI-1B2 |
| IRIS_313-9190 | ERS468308 | -0.058969 | -0.0280454 | 0.0100771 | XI-adm |
| IRIS_313-11871 | ERS469025 | -0.0588726 | -0.0386231 | -0.00119069 | XI-1A |
| CX54 | ERS470705 | -0.0586752 | -0.0399389 | 0.00484019 | XI-3B2 |
| IRIS_313-10651 | ERS469471 | -0.0581243 | -0.0336848 | 0.00106466 | XI-3B1 |
| IRIS_313-10820 | ERS469623 | -0.0576055 | -0.0463394 | 0.00188338 | XI-3A |
| CX388 | ERS470674 | -0.0572151 | -0.0477387 | 0.00283701 | XI-adm |
| CX133 | ERS470499 | -0.0570418 | -0.0459518 | -0.00068281 | XI-1A |

| | | | | | |
|---|---|---|---|---|---|
| IRIS_313-10977 | ERS469780 | -0.0570321 | -0.00487886 | 0.00585596 | XI-2A |
| IRIS_313-10239 | ERS468182 | -0.0570282 | -0.0374093 | 0.00935566 | XI-3B2 |
| IRIS_313-10990 | ERS469823 | -0.0564825 | -0.0383818 | 0.00169531 | XI-3B2 |
| CX2 | ERS470533 | -0.0564585 | -0.0523209 | 0.000578166 | XI-adm |
| B131 | ERS470341 | -0.0564321 | -0.0408457 | 0.00742544 | XI-3B2 |
| IRIS_313-8568 | ERS467875 | -0.055336 | 0.00442048 | -0.00537586 | XI-2B |
| B105 | ERS470315 | -0.0552655 | -0.043941 | 0.00320574 | XI-3B2 |
| CX145 | ERS470509 | -0.0540277 | -0.0491946 | -0.000163537 | XI-adm |
| B021 | ERS470239 | -0.0533877 | -0.0394543 | 0.00328414 | XI-3B2 |
| IRIS_313-11580 | ERS468774 | -0.0532945 | -0.0404603 | 0.00576577 | XI-3B2 |
| CX232 | ERS470556 | -0.0528192 | -0.0468444 | -0.0016278 | XI-3A |
| IRIS_313-11478 | ERS468688 | -0.0518884 | 0.0459305 | -0.0239676 | XI-2B |
| CX340 | ERS470624 | -0.0478677 | -0.051854 | -0.00424646 | XI-1B2 |
| IRIS_313-7799 | ERS468561 | -0.0444287 | 0.0492637 | -0.0292399 | XI-2B |
| CX60 | ERS470714 | -0.0422686 | -0.0414381 | 0.0125546 | XI-1B1 |
| IRIS_313-7651 | ERS468577 | -0.0393934 | 0.047613 | -0.0222178 | XI-2B |
| IRIS_313-11558 | ERS468755 | -0.0364368 | -0.019509 | -0.000923023 | XI-2A |
| IRIS_313-7758 | ERS468593 | -0.0334407 | 0.0492133 | -0.0292741 | XI-2B |
| #N/A | Hassawi_UDI0001 | -0.0271823 | 0.146839 | -0.0435627 | #N/A |
| IRIS_313-10845 | ERS469648 | -0.021389 | 0.169539 | -0.0324196 | cA2 |
| IRIS_313-10623 | ERS469461 | -0.0191007 | 0.189213 | -0.0501797 | cA2 |
| IRIS_313-11324 | ERS470116 | -0.0188249 | 0.210666 | -0.0690204 | cA1 |
| IRIS_313-11618 | ERS468796 | -0.018514 | 0.17499 | -0.0572824 | cA1 |
| IRIS_313-8321 | ERS467775 | -0.01826 | 0.178228 | -0.0325326 | cA2 |
| IRIS_313-11963 | ERS469108 | -0.018256 | 0.196954 | -0.0467947 | cA2 |
| IRIS_313-9422 | ERS467785 | -0.0176637 | 0.200251 | -0.0523136 | cA2 |
| IRIS_313-8641 | ERS467862 | -0.0174044 | 0.180574 | -0.064525 | cA1 |
| IRIS_313-12141 | ERS469232 | -0.017045 | 0.20369 | -0.0539204 | cA2 |
| IRIS_313-10965 | ERS469767 | -0.0168908 | 0.203773 | -0.0743811 | cA1 |
| IRIS_313-11059 | ERS469860 | -0.01689 | 0.230074 | -0.0762588 | cA2 |
| IRIS_313-11323 | ERS470115 | -0.0165939 | 0.197681 | -0.0600572 | cA2 |
| IRIS_313-10891 | ERS469697 | -0.0164729 | 0.116956 | -0.0497563 | cA1 |
| IRIS_313-9626 | ERS467787 | -0.0162496 | 0.22974 | -0.0728356 | cA2 |
| IRIS_313-11602 | ERS468789 | -0.0131326 | 0.199231 | -0.0645411 | cA1 |
| CX63 | ERS470716 | -0.0104227 | 0.195413 | -0.0497807 | cA1 |
| IRIS_313-11163 | ERS469965 | -0.0104153 | 0.195823 | -0.0510829 | cA2 |
| IRIS_313-11617 | ERS468795 | -0.0103709 | 0.149772 | -0.0483043 | cA1 |

| | | | | | |
|---|---|---|---|---|---|
| IRIS_313-10150 | ERS467826 | -0.00961182 | 0.185509 | -0.0693531 | cA1 |
| CX368 | ERS470653 | -0.00877361 | 0.195191 | -0.0671144 | cA1 |
| IRIS_313-10484 | ERS469374 | 0.0100683 | -0.0418826 | -0.020734 | XI-1A |
| IRIS_313-11032 | ERS469830 | 0.0418581 | 0.0918056 | 0.200327 | cB |
| IRIS_313-11625 | ERS468804 | 0.0603353 | 0.0652684 | 0.24408 | cB |
| IRIS_313-11350 | ERS470143 | 0.0737194 | 0.0551979 | 0.298018 | cB |
| IRIS_313-9682 | ERS467908 | 0.0750636 | 0.0345183 | 0.202634 | cB |
| IRIS_313-8813 | ERS467771 | 0.0786591 | 0.0559983 | 0.275588 | cB |
| CX59 | ERS470712 | 0.0793496 | 0.0635774 | 0.317507 | cB |
| IRIS_313-11567 | ERS468762 | 0.0851407 | 0.0625521 | 0.336157 | cB |
| IRIS_313-11218 | ERS470009 | 0.0854183 | 0.0578897 | 0.281858 | cB |
| CX65 | ERS470718 | 0.0914705 | 0.0673024 | 0.335119 | cB |
| CX66 | ERS470719 | 0.0922056 | 0.0667768 | 0.341754 | cB |
| IRIS_313-9301 | ERS467891 | 0.109148 | -0.0364325 | -0.0534322 | GJ-trop2 |
| IRIS_313-10080 | ERS468425 | 0.109315 | -0.0374923 | -0.041749 | GJ-trop1 |
| IRIS_313-10817 | ERS469619 | 0.111458 | -0.0343434 | -0.0469549 | GJ-trop2 |
| IRIS_313-11005 | ERS469800 | 0.111688 | -0.0321278 | -0.0398301 | GJ-trop2 |
| IRIS_313-10790 | ERS469599 | 0.112454 | -0.0337664 | -0.0491415 | GJ-trop2 |
| B071 | ERS470285 | 0.114691 | -0.0327558 | -0.0323888 | GJ-tmp |
| IRIS_313-10889 | ERS469694 | 0.114797 | -0.026388 | -0.0225572 | GJ-subtrp |
| IRIS_313-10841 | ERS469645 | 0.117049 | -0.0388164 | -0.0601148 | GJ-trop2 |
| IRIS_313-10888 | ERS469693 | 0.117324 | -0.0292782 | -0.0223761 | GJ-subtrp |
| CX129 | ERS470494 | 0.117564 | -0.0363772 | -0.0591147 | GJ-trop2 |
| CX151 | ERS470516 | 0.117624 | -0.0309357 | -0.0470096 | GJ-trop1 |
| IRIS_313-12252 | ERS469282 | 0.117661 | -0.0318472 | -0.0201454 | GJ-subtrp |
| IRIS_313-10895 | ERS469711 | 0.117829 | -0.0267513 | -0.0134183 | GJ-subtrp |
| IRIS_313-10936 | ERS469736 | 0.118409 | -0.0372002 | -0.0603893 | GJ-trop2 |
| IRIS_313-10789 | ERS469598 | 0.119008 | -0.0384674 | -0.0644515 | GJ-trop2 |
| IRIS_313-10485 | ERS469375 | 0.119404 | -0.0398132 | -0.0540498 | GJ-trop1 |
| CX248 | ERS470569 | 0.120041 | -0.0337178 | -0.0458205 | GJ-trop1 |
| IRIS_313-11183 | ERS469978 | 0.1207 | -0.0347392 | -0.0422501 | GJ-trop1 |
| B245 | ERS470443 | 0.121851 | -0.0305514 | -0.0244307 | GJ-subtrp |
| IRIS_313-12029 | ERS469163 | 0.122139 | -0.032067 | -0.0214287 | GJ-subtrp |
| IRIS_313-11007 | ERS469803 | 0.124145 | -0.0430594 | -0.0785441 | GJ-trop2 |
| IRIS_313-10805 | ERS469652 | 0.124284 | -0.0433019 | -0.0787283 | GJ-trop2 |
| IRIS_313-12350 | ERS469350 | 0.124341 | -0.0337371 | -0.0239452 | GJ-subtrp |
| IRIS_313-12262 | ERS469313 | 0.124731 | -0.0334908 | -0.0208516 | GJ-subtrp |

| | | | | | |
|---|---|---|---|---|---|
| B154 | ERS470364 | 0.125825 | -0.038879 | -0.0521342 | GJ-tmp |
| IRIS_313-12353 | ERS469353 | 0.126338 | -0.0340323 | -0.0241658 | GJ-subtrp |
| IRIS_313-8180 | ERS468521 | 0.126464 | -0.0422091 | -0.0631925 | GJ-trop1 |
| IRIS_313-11077 | ERS469880 | 0.126573 | -0.032351 | -0.0200944 | GJ-subtrp |
| IRIS_313-8111 | ERS468488 | 0.12717 | -0.0346128 | -0.0445196 | GJ-tmp |
| IRIS_313-8148 | ERS468511 | 0.127504 | -0.0362349 | -0.0456917 | GJ-tmp |
| CX132 | ERS470498 | 0.128356 | -0.0388749 | -0.0572345 | GJ-trop1 |
| CX140 | ERS470504 | 0.128532 | -0.038251 | -0.0525014 | GJ-tmp |
| IRIS_313-12018 | ERS469157 | 0.129084 | -0.0375988 | -0.0549106 | GJ-trop1 |
| IRIS_313-10258 | ERS467900 | 0.129501 | -0.0357679 | -0.0418475 | GJ-tmp |
| IRIS_313-10567 | ERS469425 | 0.129601 | -0.038598 | -0.0528003 | GJ-tmp |
| CX77 | ERS470730 | 0.129686 | -0.0425837 | -0.0645328 | GJ-trop1 |
| IRIS_313-7912 | ERS468612 | 0.129971 | -0.0384207 | -0.0623402 | GJ-trop1 |
| IRIS_313-10437 | ERS469356 | 0.13149 | -0.0401047 | -0.0550657 | GJ-tmp |
| IRIS_313-8141 | ERS468481 | 0.131521 | -0.0407819 | -0.0619746 | GJ-tmp |
| IRIS_313-10618 | ERS469483 | 0.133181 | -0.0402246 | -0.0578066 | GJ-tmp |

**Supplementary Table S5. The number of inversions between the genomes of Hassawi and the rice population reference panel.**

| Acronyms - Query genome | Query genome | Number of Inversion | Length of Inversion (bp) |
|---|---|---|---|
| IRGSP | GJ_temp.IRGSP | 241 | 13725664 |
| CM | GJ_subtrp.CHAOMEO | 223 | 9192399 |
| AZ | GJ_trop1.Azucena | 218 | 10368991 |
| KN | GJ_trop2.KETANNANGKA | 219 | 8486435 |
| ARC | cB.ARC10497 | 212 | 7748408 |
| IR64 | XI_1B1.IR64 | 165 | 6548479 |
| PR106 | XI_1B2.PR106 | 160 | 6757298 |
| ZS97 | XI_1A.ZhenShan97 | 174 | 5918719 |
| LX | XI_3B2.LIUXU | 183 | 8212805 |
| LIMA | XI_3A.LIMA | 176 | 8627439 |
| MH63 | XI_adm.MH63 | 170 | 7207335 |
| GS | XI_2A.GOBOLSAIL | 154 | 5844776 |
| KYG | XI_3B1.KHAOYAIGUANG | 171 | 6894111 |
| LM | XI_2B.LARHAMUGAD | 162 | 4846724 |
| NABO | cA2.NATELBORO | 129 | 9520888 |
| N22 | cA1.N22 | 118 | 7175315 |

**Supplementary Table S6. The number of insertions/deletions between the genomes of Hassawi and the rice population reference panel.**

| Acronyms-Query genome | Query genome | Number of Deletion | Length of Deletion | Number of Insertion | Length of Insertion (bp) |
|---|---|---|---|---|---|
| IRGSP | GJ_temp.IRGSP | 15078 | 25380373 | 11863 | 9538910 |
| CM | GJ_subtrp.CHAOMEO | 13804 | 24250320 | 12014 | 9932791 |
| AZ | GJ_trop1.Azucena | 13650 | 24234423 | 11953 | 9938190 |
| KN | GJ_trop2.KETANNANGKA | 13502 | 24004466 | 11776 | 10016093 |
| ARC | cB.ARC10497 | 12460 | 21269012 | 11273 | 9247768 |
| IR64 | XI_1B1.IR64 | 10478 | 19316935 | 9131 | 9170614 |
| PR106 | XI_1B2.PR106 | 10473 | 19044507 | 9251 | 9149829 |
| ZS97 | XI_1A.ZhenShan97 | 10349 | 20213422 | 9277 | 9400765 |
| LX | XI_3B2.LIUXU | 11535 | 19335541 | 9601 | 9323532 |
| LIMA | XI_3A.LIMA | 10889 | 20646376 | 9706 | 9532170 |
| MH63 | XI_adm.MH63 | 10640 | 19640252 | 9519 | 9053516 |
| GS | XI_2A.GOBOLSAIL | 9921 | 18340455 | 8689 | 8572444 |
| KYG | XI_3B1.KHAOYAIGUANG | 10481 | 19387433 | 9371 | 9288061 |
| LM | XI_2B.LARHAMUGAD | 11848 | 17849439 | 8918 | 8200853 |
| NABO | cA2.NATELBORO | 11011 | 14233992 | 7353 | 6058669 |
| N22 | cA1.N22 | 7296 | 12609241 | 6296 | 5492123 |

**Supplementary Table S7: Number of genes obtained by annotating the Hassawi genome.**

| | Gene # | Min. CDS length (nt) | Max CDS length (nt) | Average CDS length (nt) | Average hint support % |
|---|---|---|---|---|---|
| Chr01 | 4,832 | 201 | 13,188 | 1143.03 | 22.41 |
| Chr02 | 3,862 | 201 | 12,732 | 1160.65 | 23.15 |
| Chr03 | 4,192 | 201 | 11,487 | 1153.82 | 23.84 |
| Chr04 | 3,176 | 201 | 12,696 | 1142.49 | 21.67 |
| Chr05 | 2,880 | 201 | 7,398 | 1103.62 | 21.24 |
| Chr06 | 2,949 | 201 | 13,143 | 1114.72 | 20.71 |
| Chr07 | 2,886 | 201 | 11,037 | 1106.35 | 19.94 |
| Chr08 | 2,518 | 201 | 10,083 | 1069.15 | 19.84 |
| Chr09 | 2,110 | 201 | 13,521 | 1083.55 | 19.96 |
| Chr10 | 2,156 | 201 | 14,454 | 1090.72 | 18.15 |
| Chr11 | 2,485 | 201 | 6,168 | 1108.01 | 16.84 |
| Chr12 | 2,166 | 201 | 10,938 | 1105.52 | 18.59 |
| Subgenome1-Chr01 | 238 | 210 | 819 | 369.24 | 0 |
| Total | 36,450 | 201 | 14,454 | 1116.47 | 20.84 |

**Supplementary Table S8. Genotyping of the $T_0$ and $T_1$ of the *hd4* mutant lines.** The mutant lines are listed. Table shows the targeted gene, mutant generation, plant ID, the genotype detected compared to the wild-type sequence and the mutation type. sgRNA, PAM, and insertion sequences are written with light blue, red, and orange respectively.

| Target | Generation | Plant ID | Genotype | Mutation |
|---|---|---|---|---|
| ***Hd4*** | | WT | GGCGGCTGTTGCTCCCGCCATCGCCACGATGATGATGGATTC<br>GGCGGCTGTTGCTCCCGCCATCGCCACGATGATGATGGATTC | Wild type |
| | $T_0$ | *hd4-530* | GGCGGCTGTTGCTCCCGCCATCGCCACGATG-TGATGGATTC -1<br>GGCGGCTGTTGCTCCCGCCATCGCCACGATG-TGATGGATTC -1 | Homozygous |
| | $T_0$ | *hd4-534* | GGCGGCTGTTGCTCCCGCCATCGCCACGATG-TGATGGATTC -1<br>GGCGGCTGTTGCTCCCGCCATCGCCACGATGA-GATGGATTC -1 | Biallelic |
| | $T_0$ | *hd4-537* | GGCGGCTGTTGCTCCCGCCATCGCCACGATGATTGATGGATTC +1<br>GGCGGCTGTTGCTCCCGCCATCGCCACGATGA-GATGGATTC -1 | Biallelic |
| | $T_0$ | *hd4-540* | GGCGGCTGTTGCTCCCGCCATCGCCACGATG-TGATGGATTC -1<br>GGCGGCTGTTGCTCCCGCCATCGCCACGATGA-GATGGATTC -1 | Biallelic |
| | $T_0$ | *hd4-554* | GGCGGCTGTTGCTCCCGCCATCGCCACGATGATGATGGATTC 0<br>GGCGGCTGTTGCTCCCGCCATCGCCACGATG-TGATGGATTC -1<br>GGCGGCTGTTGCTCCCGCCATCGCCACGATGAATGATGGATTC +1 | Chimeric |
| | $T_0$ | *hd4-557* | GGCGGCTGTTGCTCCCGCCATCGCCACGATGATGATGGATTC 0<br>GGCGGCTGTTGCTCCCGCCATCGCCACGATGAATGATGGATTC +1 | Heterozygous |
| | $T_0$ | *hd4-562* | GGCGGCTGTTGCTCCCGCCATCGCCACGATG-TGATGGATTC -1<br>GGCGGCTGTTGCTCCCGCCATCGCCACGATG-TGATGGATTC -1 | Homozygous |
| | $T_0$ | *hd4-563* | GGCGGCTGTTGCTCCCGCCATCGCCACGATG-TGATGGATTC -1<br>GGCGGCTGTTGCTCCCGCCATCGCCACGATGA-GATGGATTC -1 | Biallelic |
| | $T_1$ | *hd4-530.1* | GGCGGCTGTTGCTCCCGCCATCGCCACGATG-TGATGGATTC -1<br>GGCGGCTGTTGCTCCCGCCATCGCCACGATG-TGATGGATTC -1 | Homozygous |
| | $T_1$ | *hd4-530.7* | GGCGGCTGTTGCTCCCGCCATCGCCACGATG-TGATGGATTC -1<br>GGCGGCTGTTGCTCCCGCCATCGCCACGATG-TGATGGATTC -1 | Homozygous |
| | $T_1$ | *hd4-530.15* | GGCGGCTGTTGCTCCCGCCATCGCCACGATG-TGATGGATTC -1<br>GGCGGCTGTTGCTCCCGCCATCGCCACGATG-TGATGGATTC -1 | Homozygous |

**Supplementary Table S9. Genotyping of the $T_0$ and $T_2$ of the *Mx* mutant lines.** The table shows the plant ID, the targeted gene, and the genotype detected compared to the wild-type sequence. sgRNA, PAM, and insertion sequences are written in light blue, red, and black

| Plant ID | Target | | | | | | | | | | | |
|---|---|---|---|---|---|---|---|---|---|---|---|---|
| | *Hd2* | | *Hd4* | | *Hd5* | | *GS3* | | *Gn1a* | | *Sd1* | |
| WT | CGATGACTCCACCAGGCAGGTGG | | CGCCATCGCCACGATGATGATGG | | GACGGTGCAGGAGTGCGTGTCGG | | GACGCGCTCCACCGCGAGATCGG | | GCCGCTCATCCGCGCCGACGAGG | | GGACATGCCCGTGGTCGACGTGG | |
| $T_0$ line1 | CGATGACTCCACCAGG----TGG<br>CGATGACTCCA-------GGTGG | -4<br>-7 | CGCCATCGCCACGATGATGATGG<br>CGCCATCGCCACGAT--TGATGG<br>CGCCATCGCC--------GATGG | 0<br>-2<br>-8 | GACGGTGCAGGAGTGC-TGTCGG<br>GACGGTGCAGGAGTGCGtTGTCGG | -1<br>+1 | GACGCGCTCCACCGCGAGATCGG<br>GACGCGCTCCAC----AGATCGG | 0<br>-4 | GCCGCTCATCCGCGCCGACGAGG<br>GCCGCTCATCCGCGCCGaACGAGG | 0<br>+1 | GGACATGCCCGTGGTC-ACGTGG<br>GGACATGCCCGTGGTC-ACGTGG | -1<br>-1 |
| $T_0$ line2 | CGATGACTCCACCAGG----TGG<br>CGATGACTCCA-------GGTGG | -4<br>-7 | CGCCATCGCC--------GATGG<br>CGCCATCGCC--------GATGG | -8<br>-8 | GACGGTGCAGGAGTGC-TGTCGG<br>GACGGTGCAGGAGTGCGtTGTCGG | -1<br>+1 | GACGCGCTCCAC----AGATCGG<br>GACGCGCTCCAC----AGATCGG | -4<br>-4 | GCCGCTCATCCGCGCCGACGAGG<br>GCCGCTCATCCGCGCC-ACGAGG<br>GCCGCTCATCCGCGCCGaACGAGG | 0<br>-1<br>+1 | GGACATGCCCGTGGTCGACGTGG<br>GGACATGCCCGTGGTC-ACGTGG | 0<br>-1 |
| $T_0$ line3 | CGATGACTCCACCAGG----TGG<br>CGATGACTCCACCAGG----TGG | -4<br>-4 | CGCCATCGCCACGATGATGATGG<br>CGCCATCGCCACGATGA-GATGG | 0<br>-1 | GACGGTGCAGGAGTGC-TGTCGG<br>GACGGTGCAGGAGTGCGtTGTCGG | -1<br>+1 | GACGCGCTCCACCGCGAGATCGG<br>GACGCGCTCCAC----AGATCGG | 0<br>-4 | GCCGCTCATCCGCGCCGACGAGG<br>GCCGCTCATCCGCGC--ACGAGG | 0<br>-2 | GGACATGCCCGTGGTCGACGTGG<br>GGACATGCCCGTGGTCGACGTGG | 0<br>0 |
| $T_0$ line4 | CGATGACTCCACCAGG----TGG<br>CGATGACTCCACCAGG----TGG | -4<br>-4 | CGCCATCGCCACGATGATGATGG<br>CGCCATCGCCACGATGA-GATGG | 0<br>-1 | GACGGTGCAGGAGTGCGtTGTCGG<br>GACGGTGCAGGAGTGCGtTGTCGG | +1<br>+1 | GACGCGCTCCACCGCGAGATCGG<br>GACGCGCTCCACCGCGA--TCGG | 0<br>-2 | GCCGCTCATCCGCGCCGACGAGG<br>GCCGCTCATCCGCGCCGACGAGG | 0<br>0 | GGACATGCCCGTGGTCGACGTGG<br>GGACATGCCCGTGGTCGACGTGG | 0<br>0 |
| $T_2$ line1 | CGATGACTCCACCAGG----TGG<br>CGATGACTCCACCAGG----TGG | -4<br>-4 | CGCCATCGCCACGATGATGATGG<br>CGCCATCGCCACGATGA-GATGG<br>CGCCATCGCC--------GATGG | 0<br>-1<br>-8 | GACGGTGCAGGAGTGC-TGTCGG<br>GACGGTGCAGGAGTGC-TGTCGG | -1<br>-1 | GACGCGCTCCACCGCGAGATCGG<br>GACGCGCTCCACCGCGAGATCGG | 0<br>0 | GCCGCTCATCCGCGCCGACGAGG<br>GCCGCTCATCCGCGCC-ACGAGG<br>GCCGCTCATCCGCGCCGaACGAGG | 0<br>-1<br>+1 | GGACATGCCCGTGGTCGACGTGG<br>GGACATGCCCGTGGTC-ACGTGG | 0<br>-1 |
| $T_2$ line2 | CGATGACTCCACCAGG----TGG<br>CGATGACTCCA-------GGTGG | -4<br>-7 | CGCCATCGCC--------GATGG<br>CGCCATCGCC--------GATGG | -8<br>-8 | GACGGTGCAGGAGTGC-TGTCGG<br>GACGGTGCAGGAGTGC-TGTCGG | -1<br>-1 | GACGCGCTCCACCGCGAGATCGG<br>GACGCGCTCCACCGCGAGATCGG | 0<br>0 | GCCGCTCATCCGCGCC-ACGAGG<br>GCCGCTCATCCGCGCCGaACGAGG | -1<br>+1 | GGACATGCCCGTGGTCGACGTGG<br>GGACATGCCCGTGGTC-ACGTGG | 0<br>-1 |
| $T_2$ line3 | CGATGACTCCACCAGG----TGG<br>CGATGACTCCACCAGG----TGG | -4<br>-4 | CGCCATCGCCACGATGA-GATGG<br>CGCCATCGCCACGATGA-GATGG | -1<br>-1 | GACGGTGCAGGAGTGC-TGTCGG<br>GACGGTGCAGGAGTGC-TGTCGG | -1<br>-1 | GACGCGCTCCACCGCGAGATCGG<br>GACGCGCTCCACCGCGAGATCGG | 0<br>0 | GCCGCTCATCCGCGCC-ACGAGG<br>GCCGCTCATCCGCGCCGaACGAGG | -1<br>+1 | GGACATGCCCGTGGTCGACGTGG<br>GGACATGCCCGTGGTCGACGTGG | 0<br>0 |
| $T_2$ line4 | CGATGACTCCA-------GGTGG<br>CGATGACTCCA-------GGTGG | -7<br>-7 | CGCCATCGCCACGATGATGATGG<br>CGCCATCGCCACGATGA-GATGG<br>CGCCATCGCC--------GATGG | 0<br>-1<br>-8 | GACGGTGCAGGAGTGC-TGTCGG<br>GACGGTGCAGGAGTGC-TGTCGG | -1<br>-1 | GACGCGCTCCACCGCGAGATCGG<br>GACGCGCTCCAC----AGATCGG | 0<br>-4 | GCCGCTCATCCGCGCCGACGAGG<br>GCCGCTCATCCGCGCCGACGAGG | 0<br>0 | GGACATGCCCGTGGTCGACGTGG<br>GGACATGCCCGTGGTCGACGTGG | 0<br>0 |
| $T_2$ line5 | CGATGACTCCACCAGG----TGG<br>CGATGACTCCA-------GGTGG | -4<br>-7 | CGCCATCGCC--------GATGG<br>CGCCATCGCC--------GATGG | -8<br>-8 | GACGGTGCAGGAGTGC-TGTCGG<br>GACGGTGCAGGAGTGC-TGTCGG | -1<br>-1 | GACGCGCTCCAC----AGATCGG<br>GACGCGCTCCAC----AGATCGG | -4<br>-4 | GCCGCTCATCCGCGCCGaACGAGG<br>GCCGCTCATCCGCGCCGaACGAGG | +1<br>+1 | GGACATGCCCGTGGTCGACGTGG<br>GGACATGCCCGTGGTCGACGTGG | 0<br>0 |
| $T_2$ line6 | CGATGACTCCA-------GGTGG<br>CGATGACTCCA-------GGTGG | -7<br>-7 | CGCCATCGCC--------GATGG<br>CGCCATCGCC--------GATGG | -8<br>-8 | GACGGTGCAGGAGTGC-TGTCGG<br>GACGGTGCAGGAGTGC-TGTCGG | -1<br>-1 | GACGCGCTCCACCGCGAGATCGG<br>GACGCGCTCCAC----AGATCGG | 0<br>-4 | GCCGCTCATCCGCGCCGaACGAGG<br>GCCGCTCATCCGCGCCGaACGAGG | +1<br>+1 | GGACATGCCCGTGGTCGACGTGG<br>GGACATGCCCGTGGTCGACGTGG | 0<br>0 |
| $T_2$-*Mx* | CGATGACTCCA-------GGTGG<br>CGATGACTCCA-------GGTGG | -7<br>-7 | CGCCATCGCCACGAT--TGATGG<br>CGCCATCGCCACGAT--TGATGG | -2<br>-2 | GACGGTGCAGGAGTGC-TGTCGG<br>GACGGTGCAGGAGTGC-TGTCGG | -1<br>-1 | GACGCGCTCCAC----AGATCGG<br>GACGCGCTCCAC----AGATCGG | -4<br>-4 | GCCGCTCATCCGCGCCGaACGAGG<br>GCCGCTCATCCGCGCCGaACGAGG | +1<br>+1 | GGACATGCCCGTGGTCGACGTGG<br>GGACATGCCCGTGGTC-ACGTGG | 0<br>-1 |

respectively.

**Supplementary Table S10.** Fold-change table of all compounds significantly increased by more than three-fold in one of the modified lines.

| Super Pathway | Sub Pathway | Biochemical Name | RR_MX / RR_WT | RR_HD4 / RR_WT |
|---|---|---|---|---|
| Amino Acid | Aromatic amino acid metabolism (PEP derived) | tryptophan | 5.72 | 1.86 |
| | | tryptamine | 2.32 | 3.49 |
| | | tyramine | 4.89 | 2.67 |
| | Glutamate family (alpha-ketoglutarate derived) | glutamine | 4.42 | 2.83 |
| | Glutathione metabolism | glutathione, reduced (GSH) | 1.91 | 3.88 |
| | | cysteine-glutathione disulfide | 1.28 | 3.81 |
| | | cysteinylglycine | 1.42 | 3.09 |
| Lipids | Choline metabolism | choline phosphate | 2.35 | 3.4 |
| | | 2-dimethylaminoethanol | 2.66 | 4.99 |
| Cofactors, Prosthetic Groups, Electron Carriers | Nicotinate and nicotinamide metabolism | nicotinamide riboside | 3.89 | 5.33 |
| | Thiamine metabolism | thiamin (vitamin B1) | 1.95 | 3.06 |
| | Chlorophyll and heme metabolism | pheophorbide A | 11.49 | 18.11 |
| | | pheophytin A | 15.76 | 12.51 |
| Hormone metabolism | Auxin metabolism | 2-oxindole-3-acetate | 9.35 | 1 |
| Secondary metabolism | Flavonoids | catechin | 3.05 | 5.19 |
| | Phenylpropanoids | p-coumaroylserotonin | 12.51 | 15.66 |
| | Terpenoids | carotene diol (1) | 10.16 | 11.77 |
| | | carotene diol (2) | 13.5 | 12.99 |
